# Leveraging pleiotropy to discover and interpret GWAS results for sleep-associated traits

**DOI:** 10.1101/832162

**Authors:** Sung Chun, Sebastian Akle, Athanasios Teodosiadis, Brian E. Cade, Heming Wang, Tamar Sofer, Daniel S. Evans, Katie L. Stone, Sina A. Gharib, Sutapa Mukherjee, Lyle J Palmer, David Hillman, Jerome I. Rotter, Craig L. Hanis, John A. Stamatoyannopoulos, Susan Redline, Chris Cotsapas, Shamil R. Sunyaev

**Author notes:** These authors contributed equally to this work (Sung Chun, Sebastian Akle). These authors contributed equally to this work (Chris Cotsapas, Shamil R. Sunyaev).

## Abstract

Genetic association studies of many heritable traits resulting from physiological testing often have modest sample sizes due to the cost and burden of the required phenotyping. This reduces statistical power and limits discovery of multiple genetic associations. We present a strategy to leverage pleiotropy between traits to both discover new loci and to provide mechanistic hypotheses of the underlying pathophysiology. Specifically, we combine a colocalization test with a locus-level test of pleiotropy. In simulations, we show that this approach is highly selective for identifying true pleiotropy driven by the same causative variant, thereby improves the chance to replicate the associations in underpowered validation cohorts and leads to higher interpretability. Here, as an exemplar, we use Obstructive Sleep Apnea (OSA), a common disorder diagnosed using overnight multi-channel physiological testing. We leverage pleiotropy with relevant cellular and cardio-metabolic phenotypes and gene expression traits to map new risk loci in an underpowered OSA GWAS. We identify several pleiotropic loci harboring suggestive associations to OSA and genome-wide significant associations to other traits, and show that their OSA association replicates in independent cohorts of diverse ancestries. By investigating pleiotropic loci, our strategy allows proposing new hypotheses about OSA pathobiology across many physiological layers. For example, we identify and replicate the pleiotropy across the plateletcrit, OSA and an eQTL of DNA primase subunit 1 (*PRIM1*) in immune cells. We find suggestive links between OSA, a measure of lung function (FEV_1_/FVC), and an eQTL of matrix metallopeptidase 15 (*MMP15*) in lung tissue. We also link a previously known genome-wide significant peak for OSA in the hexokinase 1 (*HK1*) locus to hematocrit and other red blood cell related traits. Thus, the analysis of pleiotropic associations has the potential to assemble diverse phenotypes into a chain of mechanistic hypotheses that provide insight into the pathogenesis of complex human diseases.

**Author Summary:** Large genetic studies with hundreds of thousands of patients have been successful at finding genetic variants that associate with disease traits in humans. However, smaller-scale studies can often have inadequate power to discover new genetic associations. Here, we use a small genetic study of Obstructive Sleep Apnea (OSA), to introduce a strategy that both helps find genetic associations and proposes biological hypotheses for the mechanisms behind those associations. To achieve this, we use large genetic studies carried out on traits that are related to OSA, and look for genetic variants that affect both OSA in our small study and the trait in question in the large study. By linking two or more traits at select loci, we were able to, among other results, find a locus that affects the expression of a gene in immune cells (DNA primase subunit 1), a marker of thrombotic and inflammatory processes (plateletcrit) and OSA. This results in a novel genetic association to OSA and a corresponding biological hypothesis behind its effect on OSA.

## Introduction

Genome-wide association studies of human phenotypes ranging from gene expression to human diseases are now routine. Cumulatively, the data indicate that complex traits are highly polygenic [1,2], and genetic correlation between these traits indicates abundant pleiotropy [3–5]. Interpreting the plethora of results raises two major challenges: first, generating testable mechanistic hypotheses about the underlying pathophysiology; and second, increasing statistical power to identify associations in studies of traits with small or moderate sample sizes. Leveraging pleiotropy can help address both of these challenges. Previous work has demonstrated that including many correlated traits in association studies increases power to detect associations common to multiple traits [3,6,7]. This approach is largely untried in genetic investigations of Obstructive Sleep Apnea (OSA). Here, we demonstrate that using shared associations between correlated traits can identify effects in under-powered studies of OSA, and that leveraging molecular and physiological endophenotypes in this way also generates clear and testable biological hypotheses.

OSA is characterized by recurrent episodes of partial or complete obstruction of the pharyngeal airway resulting in multiple physiological disturbances, including sympathetic nervous system activation, increased energy cost of breathing, intermittent hypoxemia, and wide swings in intrathoracic pressure. This disorder is highly prevalent in the general population, affecting more than 10% of middle-aged adults, with increased prevalence observed with aging, obesity, and cardiometabolic disease, and is more common in men [8]. OSA leads to sleep disruption, particularly increased sleep fragmentation and decreased proportion of restorative stages of sleep, resulting in daytime sleepiness, impaired quality of life and cognitive deficits [9].

Moreover, OSA is associated with increased rates of hypertension, incident heart disease, stroke, diabetes, depression, certain cancers, and overall mortality [10–19]. Despite the large number of epidemiological studies indicating that OSA is closely associated with these outcomes, there appear to be subgroup differences in susceptibility, e.g., middle-aged individuals and men are more likely to experience OSA-related cardiovascular disease in some studies than older individuals and women, respectively [20]. This underscores gaps in our knowledge of the pathophysiological pathways linking OSA to other diseases [21,22].

Pathophysiological pathways linking OSA to other diseases and factors that influence individual differences in susceptibility are poorly understood. While there are several effective treatments for OSA, including Continuous Positive Airway Pressure (CPAP), there appears to be substantial variation in overall clinical response and attenuation of cardiometabolic consequences, suggesting heterogeneity in both the etiology of the disease and susceptibility to its physiological disturbances.

Indices of OSA, including the Apnea-Hypopnea Index (AHI; the number of breathing pauses per hour of sleep), apnea event duration, indices of overnight hypoxemia, habitual snoring, and excessive daytime sleepiness, show substantial heritability in family studies [23]. Past studies have identified only a handful of associations with a variety of OSA-related traits. We have previously described a GWAS of OSA traits measured by overnight polysomnography in multi-ethnic cohorts totaling ∼20,000 individuals [24]. In that study, we found two genome-wide significant multiethnic associations: variants in a locus on 10q22 were associated with indices of average and minimum SpO_2_ and percentage of sleep with SpO_2_ < 90%, and variants in a locus on 2q12 were associated with minimum oxygen saturation (SpO_2_). In another study, we identified a locus in 17p11 with a male-specific effect on AHI [25]. Furthermore, in an admixture mapping study in Hispanic/Latino Americans, we identified a locus on 2q37 associated with AHI and one in a locus on 18q21 associated with AHI and SpO_2_ < 90% [26].

The low number of genetic associations reported to date only explains a small fraction of OSA trait heritability. This relative paucity of findings is driven primarily by modest sample sizes, a reflection of the expense and complexity of measuring physiological phenotypes by overnight polysomnography or respiratory polygraphy. This also limits our ability to fine-map associations down to causative variants and thus identify relevant genes. Data on hundreds of thousands of individuals – sample sizes at which GWAS designs are well-powered to detect tens of loci and, in combination with additional experiments, fine-map some of them – have yet to be collected for OSA traits based on overnight polysomnography or respiratory polygraphy and may never be available [1,2]. Biological interpretation of available genetic associations is further complicated by the observation that most GWAS effects localize to enhancer regions and other regulatory elements and are often distal to physiologically relevant genes [27,28].

Now that GWAS of massive sample sizes have been accumulated for various comorbid conditions and endophenotypes related to OSA, we hypothesize that analysis of shared associations across correlated traits can identify effects in underpowered studies of OSA and generate clear and testable biological hypotheses. A number of computational methods increase power for discovering genetic associations by capitalizing on pleiotropy between disease phenotypes or between a disease and a molecular trait such as gene expression. One common approach takes advantage of the genetic correlation among phenotypes [6,29–31].

This class of methods gains substantial additional power by pooling association signals across traits. However, such methods suffer from power loss when the correlation of genetic effect sizes is highly variable across the genome or limited to a subset of loci. For example, an approach such as MTAG is not suitable for small GWAS studies when the assumption of the homogeneity of genetic correlation is violated. The estimated genetic effect of an underpowered trait can be inflated by the strong genetic signals of well-powered traits that are pooled together even if the variant is not causative for the underpowered trait [6]. An alternative approach analyzes individual loci to detect pleiotropic alleles, with no regard to genetic correlation [4,7,32–41]. Only a handful of existing methods account for the possibility that the apparent pleiotropy is driven by the linkage disequilibrium (LD) between two distinct causative variants each of which drives only one phenotype [4,36–40].

In this study, we apply a colocalization method to detect shared associations between OSA and other related traits. We elected Joint Likelihood Mapping method (JLIM) [39] and eCAVIAR [38] as colocalization methods, but our general strategy does not depend on any specific method. We focus on a set of well powered intermediate traits which have previously been implicated in the pathobiology of OSA. Given prior GWAS studies suggesting the involvement of inflammatory genes in OSA [42–44], and cohort studies reporting high levels of inflammation, including elevations in neutrophils and monocytes in OSA [45,46], we included leukocyte and platelet related traits in our pleiotropic comparisons. Similarly, we also included red blood cell related traits given prior GWAS implicating iron metabolism [26] and erythrocyte function [24]. We will refer to these as clinical traits. In addition, OSA is associated with lung [47,48], obesity and cardiovascular-related disorders [46,49–51], and we have included clinical traits that reflect these overlaps, together with gene expression traits in tissues implicated in these diseases.

By linking different clinical and gene expression traits to OSA at specific loci, our analysis suggests new hypotheses about OSA pathobiology across many physiological layers as well as finding a new association.

## Results

### Creating a framework to identify associations in underpowered GWAS through pleiotropy

We used a colocalization method to identify pleiotropic loci, where a genetic effect drives association to two traits. First, we selected genome-wide significant loci (association *p* < 5 × 10^-8^) in our well-powered trait (here, a clinical trait), and from these we selected the subsets which also show nominal association to OSA traits (*p* < 0.01 for any SNP in the locus). We then used colocalization to directly evaluate if the association to the two traits was consistent with the same underlying effect, indicating a pleiotropic effect. This two-step strategy allowed us to distinguish between cases where there was association only in the clinical trait; where there was a shared association in both traits; and if there were distinct associations in both traits stemming from different underlying effects (Figures S1 and S2).

### Simulations

We first used simulations to assess our strategy to identify true associations in an underpowered study by detecting pleiotropic associations with a better-powered study of another trait and following up with replication in an independent cohort. To assess sensitivity and specificity, we simulated variable proportions of shared and distinct causal effects, whilst assuming high polygenicity for each trait and only one causative variant per locus. We simulated GWAS statistics for pairs of traits corresponding to our own study design: we simulated situations where no variant is causative for an underpowered trait (H_0_), where the same variant is causative for the association to both traits (H_1_), and where two distinct variants in LD drive each of the associations at a locus (H_2_) (Methods). To simulate summary statistics from a well-powered GWAS (representing our clinical traits), we sampled values from a multivariate normal (MVN) distribution using the local LD matrix as a variance-covariance parameter [52] with clinical trait sample size of 150,000. To simulate a GWAS of limited power (representing our OSA discovery cohorts), we used genotypes from ancestry-matched samples as a reference from which we simulated quantitative traits in 10,000 samples as surrogates for the OSA GWAS. Most overlapping associations are expected to be due to distinct variants [39]; we simulated scenarios with just 5% and 20% of true associations being driven by the same variant in both traits (H_1_). For example, in the first case, we simulated 2,500 loci with 3.5% corresponding to H_1_, 66.5% corresponding to H_2_ and 30% corresponding to no association in the underpowered trait (H_0_). The focus of our simulation study was the ability to replicate the association signal for the primary phenotype in an independent cohort. This means successfully discriminating between (H_1_+H_2_) and H_0_. We leave the interpretability question (discriminating H_1_ from H_2_) to Discussion.

We tested two colocalization methods – JLIM and eCAVIAR (a popular Bayesian colocalization test) [38,39]. We also tested a popular pleiotropy-informed test, conditional false discovery rate (cFDR) [32,33]. cFDR is not based on the LD-informed colocalization of the association signals. All methods appear underpowered in the discovery dataset at a sample size typical of current genetic studies of OSA (n=10,000); the methods are only able to find associations if the observed effect is larger than the true effect (Figures S3 and S4A). Among the true associations identified by JLIM, eCAVIAR and cFDR, there is an enrichment of H_1_ loci. In spite of this enrichment, many loci identified in the discovery cohort are driven by distinct variants (H_2_): (for the simulation with 5% of the true associations being H_1_, 54.7, 55.8 and 61.5% of loci identified by JLIM, eCAVIAR and cFDR, respectively, are H_2_). At the same time, almost all proportions of successfully replicated associations are driven by the same variant (H_1_) in our simulation regimes (Figures 1C-D and S4B-F).

**Figure 1.**
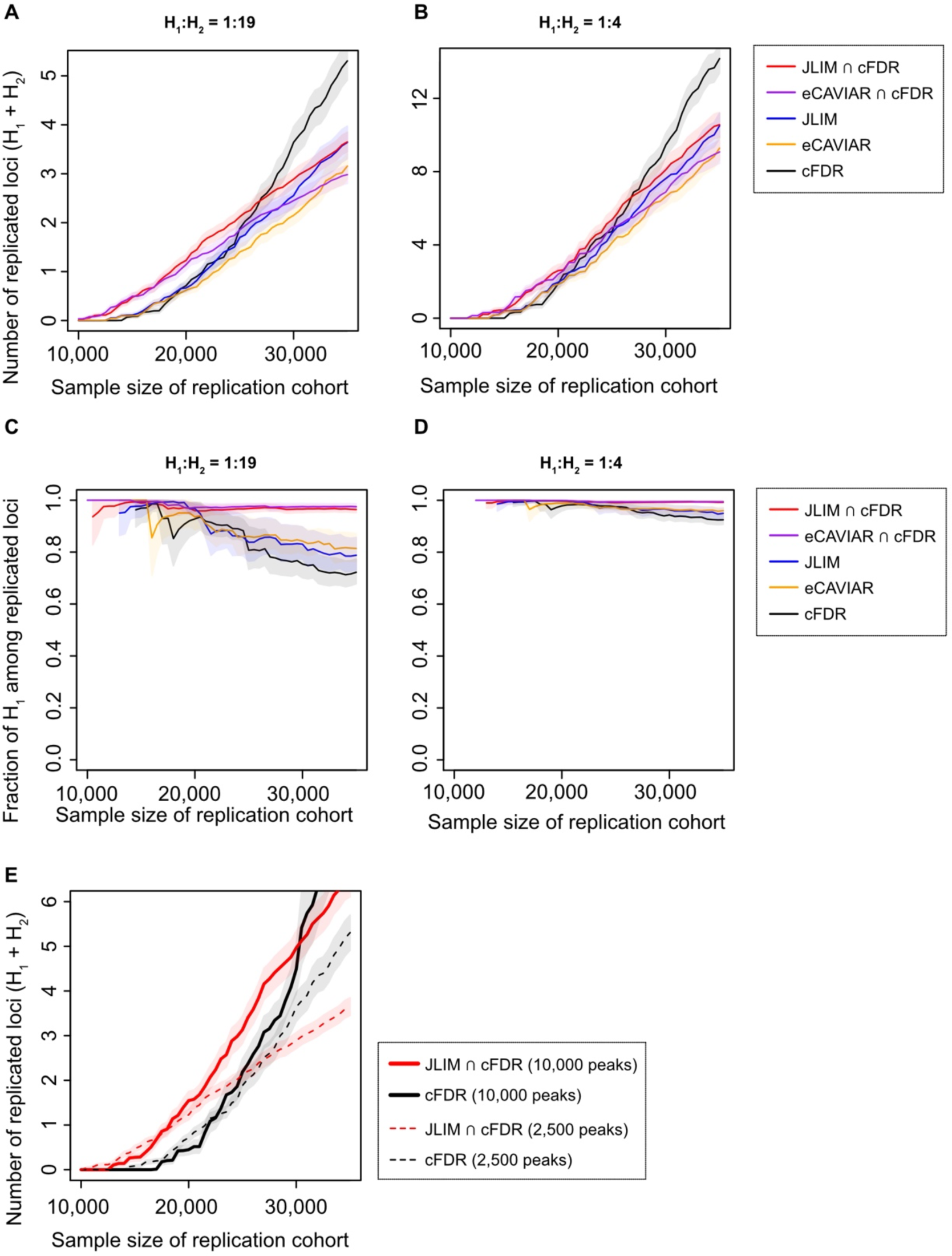
The projected number of replicated loci and proportion of shared causative variants among the replicated loci. A total of 2,500 association peaks from well-powered GWAS studies (n=150,000) were tested for the shared effect in simulated discovery cohorts (n=10,000), and then select loci were tested for replication in simulated validation cohorts of the same genetic ancestry (n=10,000-35,000). The candidate loci were identified by conditional false discovery rate (cFDR), colocalization tests (JLIM and eCAVIAR), or the intersection of cFDR and colocalization test. The P-value cutoff was set to 0.01 for cFDR and JLIM, and for eCAVIAR, the equivalent posterior threshold was calibrated using H_0_ simulation data. The 2,500 GWAS peaks consist of the loci simulating no causal effect for underpowered traits (H_0_) and those simulating the same causal effect between two traits (H_1_) or distinct causal effects (H_2_). The H_1_:H_2_ ratio was set to 1:19 (**A** and **C**) or 1:4 (**B** and **D**). The effect sizes of causative variants are correlated (*ρ* = 0.7) under H_1_ but uncorrelated under H_2_. Bonferroni correction was applied on replication tests. In Panel E, we contrast the number of replicated loci expected in Panel **A** (dashed line) with a more extreme scenario (10,000 GWAS peaks, consisting of 2,500 with H_1_:H_2_ of 1:19 and 7,500 with H_1_:H_2_ of 1:39; solid lines). In all, the proportion of H_0_ was set to 30% of all examined loci. The shaded area denotes the 95% CIs.

Because the colocalization tests (such as JLIM and eCAVIAR) and cFDR are using different features of the data, we found that taking a consensus between them can effectively exclude the majority of H_2_ from the analysis (Figure S3). In the intersection of JLIM and cFDR, only 3.5% of identified loci are H_2_; in the intersection of eCAVIAR and cFDR, only 5.3% are H_2_ (Figure S4A). Even though the elimination of H_2_ loci increases the false discovery rate (fraction of H_0_) in the discovery sample, it substantially increases the rate of successful replication. The main reason for the increase in the rate of successful replication is the prioritization of H_1_ loci of larger true effect size and the corresponding reduction of multiple testing burden at the replication stage. The benefit of this approach increases with the fraction of loci driven by distinct effects (H_2_) (Figure S5). It also increased when the fraction of loci of no true effect (H_0_) is low (as in the case of very high polygenicity [53] (Figure S6)).

To test if the consensus method allows for more replicated loci simply due to the smaller number of candidate loci, we tightened the p-value threshold of cFDR to select the same number of candidates in discovery as the consensus method (Figure S7) or to control the rate of H_0_ loci (Figure S8). We also explored a range of other simulation conditions, including varying the total number of examined well-powered trait-associated loci, correlation of effect sizes between well-powered and underpowered traits, trans-ethnic validation cohort, and the presence of multiple causative variants in locus (Figures S9-12). Consistently, the consensus method resulted in more replicated loci than cFDR alone when the sample size is limited.

Next, we compared the rate of successful replication between the consensus approach and Bayesian meta-analysis methods. In contrast to colocalization methods, the Bayesian meta-analysis approach leverages the correlation of effect sizes between traits rather than SNP-level colocalization of causative variants [29–31]. We chose to compare two such Bayesian meta-analysis methods, MetABF and CPBayes, with the consensus approach using the simulated dataset. Again, when the sample size was limited (n < 20,000), our consensus method found more replicated associations than meta-analysis approaches (Figures S13-14). As the sample size of the validation cohort increases, both MetABF and CPBayes identified more H_2_ loci, which led to more replicated loci than the consensus method. Since the meta-analysis methods are designed for a pleiotropy analysis of more than two traits, in principle, they can boost the power by using more than one well-powered trait. To test if the multi-trait meta-analysis can outperform the pairwise meta-analysis, we simulated nine additional well-powered traits, generating GWAS statistics for one underpowered trait and ten well-powered traits in each locus (Methods). For the additional well-powered traits, we randomly decided whether the simulated causative variant was to be the same as or distinct from that of the main well-powered trait. Using this simulated dataset, we examined the power of the above Bayesian methods and non-parametric meta-analysis (iGWAS) [7] in multi-trait and pairwise settings. Overall, we could not see a clear advantage of multi-trait meta-analyses over pairwise tests in our simulated data (Figure S15). Rather, particularly for CPBayes and iGWAS, multi-trait tests substantially underperformed pairwise tests. This result demonstrates that for underpowered studies, the consensus approach focused on SNP-level colocalization outperforms meta-analysis techniques in pairwise and multi-trait settings when the association signals are driven by distinct rather than the same effect in a large fraction of loci.

Finally, we found that looking at more trait combinations with more GWAS peaks results in proportionally more discoveries (Figure 1E). Overall, our simulations demonstrate that casting a wide net across many traits (increasing the number of GWAS peaks) and taking the consensus between pleiotropy mapping methods is a viable strategy to increase discoveries in under-powered studies. We therefore felt justified in pursuing this strategy using real data, to make additional discoveries in traits related to OSA.

### Identifying pleiotropic associations between clinical traits and sleep apnea-related traits

Based on clinical relevance [54] and heritability [23], we focused on four OSA-related traits measured in five European-ancestry cohorts: the apnea-hypopnea index (AHI) [25], average respiratory event (apneas or hypopneas) duration [26], and minimum and average oxygen saturation (SpO_2_) during sleep [24]. We used summary statistics from the remaining multi-ethnic cohorts in our replication effort.

We assembled a collection of GWAS summary statistics for a total of 55 candidate intermediate traits from across these physiological areas: erythroid, leukocyte and platelet counts and function, from a study combining the UK Biobank and INTERVAL datasets (170,000 individuals of European ancestry) [55]; cardiovascular, metabolic and respiratory traits from the UK Biobank (380-450,000 European ancestry participants) [56,57], and cardio-metabolic traits (36,000 European ancestry participants) [58]. We then compared associations in each of these clinical traits to our OSA traits (6,781 European ancestry participants; Table S1), to identify potential associations in the latter. A complete list of clinical traits we considered is presented in Table S2.

We tested for directional causal effects of the selected clinical traits on our OSA related traits using Mendelian Randomization (MR) [59]. Due to the low sample sizes in OSA traits, no comparison reached statistical significance after multiple test correction (Table S3). After excluding the extended MHC region and the sex chromosomes, we identified 3,191 genome-wide significant associations (*p* < 5 × 10^-8^) in the 55 clinical traits, of which 2,939 had a corresponding suggestive association to one of the four OSA traits (*p* < 0.01 at any SNP in the locus; Table S4). We then explicitly tested for evidence of pleiotropy between clinical and OSA traits using JLIM (2,142 to 2,236 tests for each OSA trait). We found evidence that in 61/2,939 of these regions the OSA and clinical trait associations are consistent with a shared, pleiotropic underlying causative variant by JLIM (false discovery rate (FDR) < 0.20) and at the same time show evidence of association to OSA by cFDR (Table 1, Table S5).

**Table 1.**
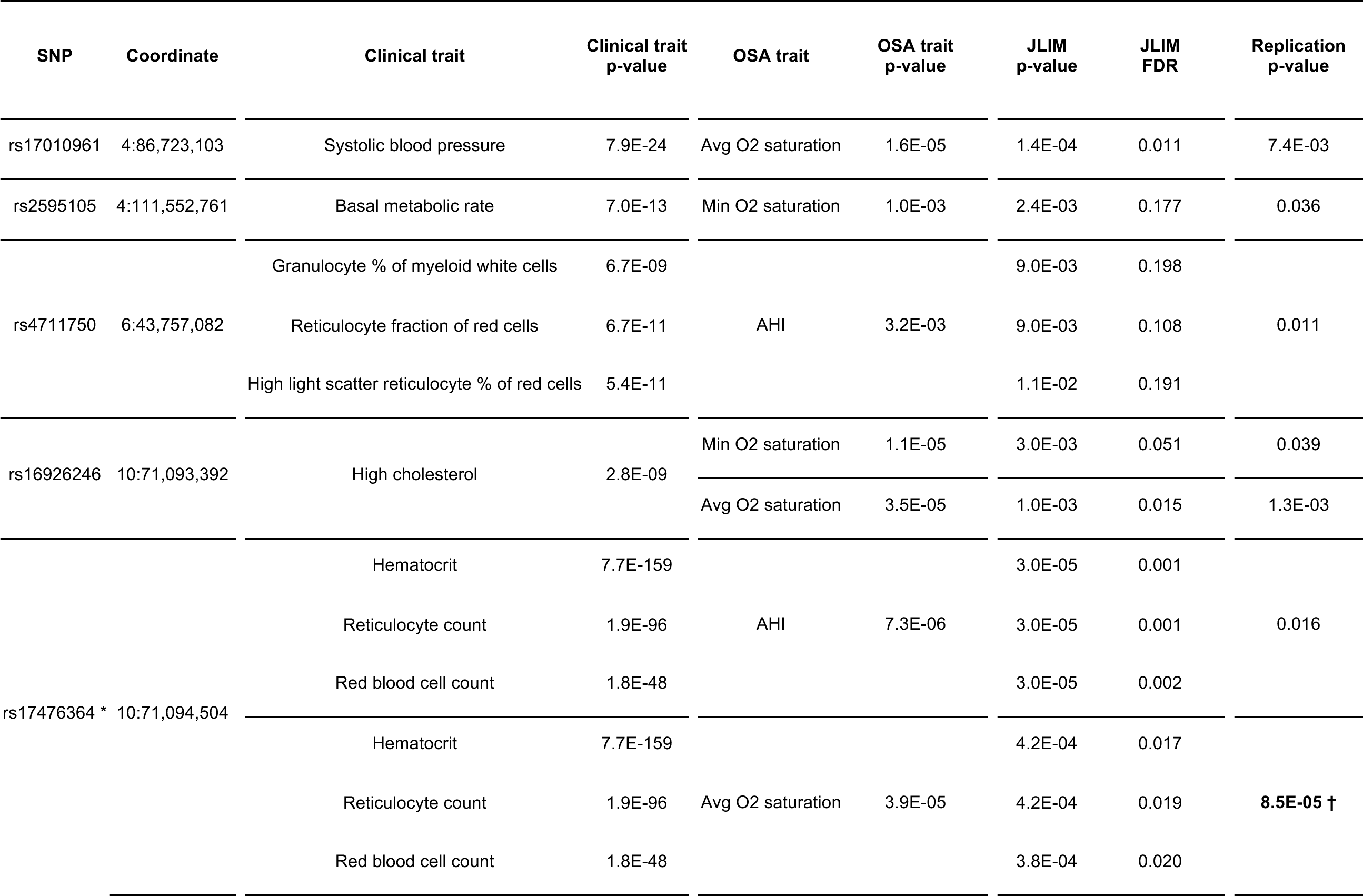
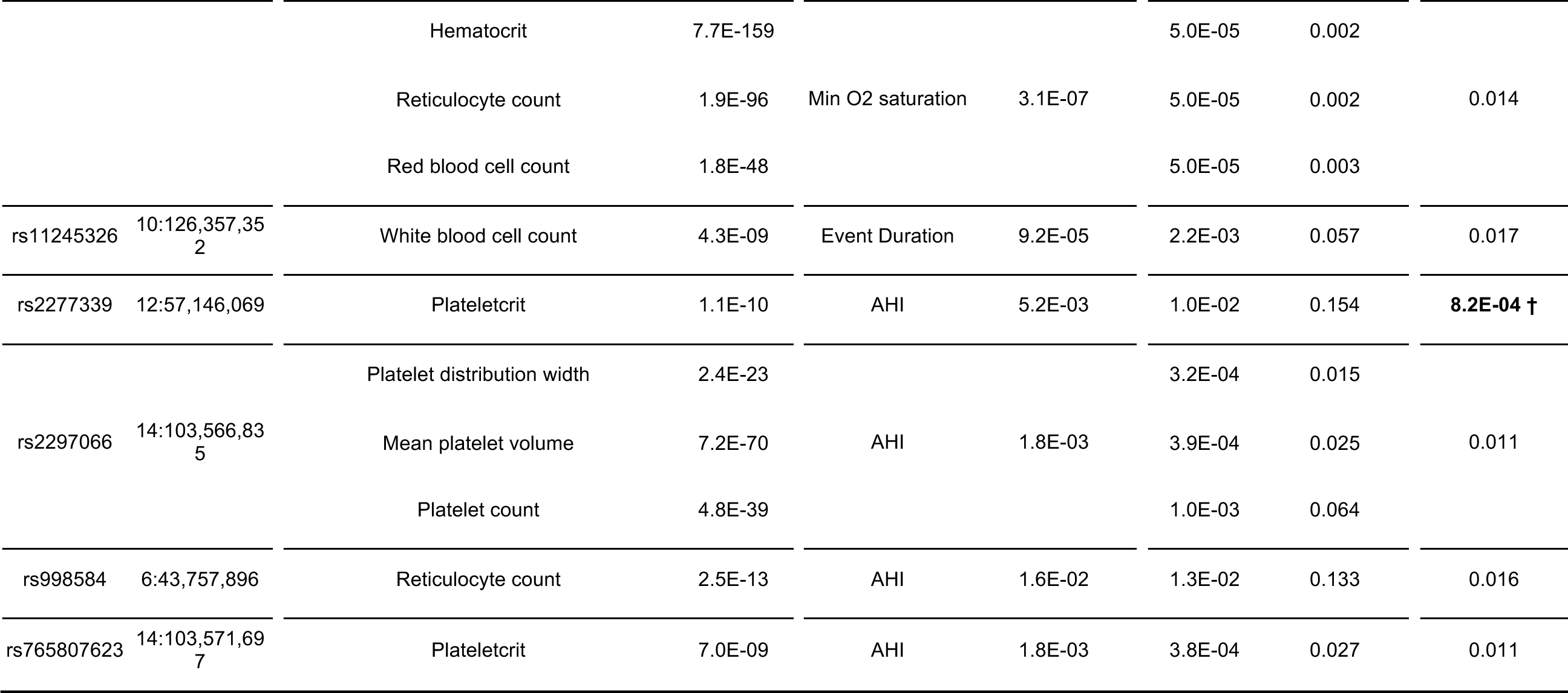
Loci with significant pleiotropic associations between a clinical trait and an OSA trait with nominally significant replication. Each row denotes one SNP and its corresponding associations to clinical and OSA traits. Each SNP may be associated with more than one clinical trait. In SNPs marked with *, not all clinical traits are shown for clarity. Here, we only show variants with nominally significant replication p-values. See Table S5 for the full table of 61 loci (137 trait pairs), including those with insignificant replication p-values. Two variants marked with † indicates pleiotropic loci with significant out of sample replication p-values after Bonferroni correction (0.05/61). AHI stands for Apnea-hypopnea index. Coordinates correspond to hg19. The clinical trait p-value column refers to the association p-value of the SNP to the clinical trait. The OSA trait p-value refers to the association p-value of the SNP to the OSA trait in meta-analyzed discovery cohorts (Table S1). The replication p-value refers to the association p-value of the SNP to the OSA trait in meta-analyzed validation cohorts (Table S6). The JLIM p-value corresponds to the tests for the shared effect between the clinical and OSA phenotypes.

To independently validate our 61 putative OSA trait associations from the discovery stage, we compiled summary statistics for the same traits in 15,594 individuals of Asian, African, European and Hispanic ancestries/backgrounds (Table S6). These individuals do not overlap with those from the cohorts used in our colocalization analysis using JLIM. We did not attempt to replicate pleiotropic associations; we only replicated the OSA association statistics. JLIM relies on local LD patterns being preserved between clinical and OSA trait cohorts, so we cannot use multi-ethnic data in our discovery analysis of pleiotropy. We found that 2/61 variants in Table S5 show significant association with the same OSA trait as the initial observation, after Bonferroni correction for the number of tests performed. The variant in SNP rs17476364 (Figure 2) links 11 of 12 red blood cell related clinical traits analyzed with average SpO_2_ during sleep. It is an intronic variant in the hexokinase 1 (*HK1*) region in chromosome 10, and has been previously reported, as it reached genome-wide significance in association to minimum and average SpO_2_ [24]. Another variant in SNP rs2277339 is a missense coding variant in DNA primase subunit 1 (*PRIM1*) in chromosome 12. It links plateletcrit (ratio of platelet volume to whole blood volume, a marker of thrombotic as well as a broad range of inflammatory processes [60]) to AHI. In the UK Biobank, it has documented significant associations to height, waist to hip-ratio, age at menopause and multiple red blood cell related traits [61]. A further 8 variants, shown in Table 1, were below nominal association *p* values < 0.05 in the replication set, but did not survive multiple test correction.

**Figure 2.**
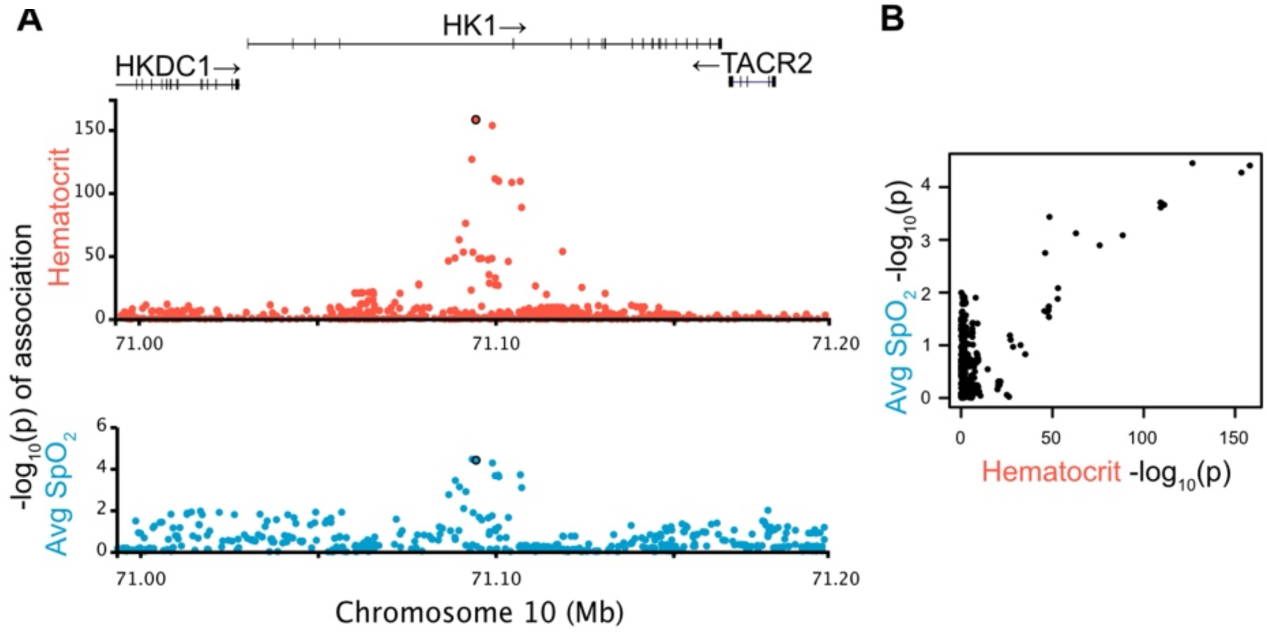
Pleiotropy at HK1 locus. **A)** Putatively pleiotropic locus linking a clinical trait (hematocrit) with an OSA trait (average SpO_2_). The evidence of shared effect between the clinical and OSA traits is significant (JLIM p = 4.2 × 10^-4^, FDR = 0.017). This SpO_2_ association replicates in validation cohorts after Bonferroni correction of 61 tests (p = 8.5 × 10^-5^). **B)** Pairwise comparison of –log10(p-values) between two traits confirms that the association signals are driven by the shared underlying effect. Each dot represents a SNP in the tested locus.

### Incorporating gene expression to construct molecular hypotheses of sleep apnea physiology

Non-coding regions with evidence of gene regulatory activity carry a large proportion of heritability in most traits analyzed in large GWAS [62]. We reasoned that some OSA causative variants would reside in such regulatory regions, and thus act on gene regulation. We therefore sought shared associations between gene expression traits and the clinical traits for which we identified a pleiotropic association in the 61 loci in Table 1 and Table S5. To do so, we compiled expression quantitative trait loci (eQTL) data for protein-coding genes expressed in lung, liver, spleen and skeletal muscle from individuals with European ancestry from the GTEx Project [63], and monocyte, T cell and neutrophil populations in individuals from BLUEPRINT [64]. We chose these tissues for potential relevance to OSA pathology: the lung is involved in OSA-related hypoxemia [47,48]; previous GWAS associations have implicated the neuromuscular junction in overnight SpO_2_ levels, and abnormalities in upper airway muscle function are fundamental mechanisms for sleep apnea [24,65]; the spleen and liver are known to mediate filtration of erythrocytes, iron homeostasis and production of inflammatory cytokines; and leukocytes are key modulators of inflammation, an antecedent risk factor of OSA development [66]. We calculated FDR based on JLIM p-values over the 5,860 comparisons against 1,009 protein-coding genes of which transcription start sites are within 1 Mb from the index SNPs of clinical traits.

We were able to identify shared associations between eQTL and clinical traits in 7/61 loci (FDR < 0.05; Table 2). This includes several notable examples, including rs2277339, one of our two replicated SNPs (Table 1). The rs2277339 SNP is a missense variant of *PRIM1* but also an eQTL for *PRIM1* levels in monocytes and T cells (Figure 3A). Another example is a locus on chromosome 16 where we find that an eQTL for *MMP15* (matrix metallopeptidase 15) expression in lung tissue is pleiotropic with a measure of lung function (FEV_1_/FVC, the ratio of forced expired volume per second to forced vital capacity), which in turn is pleiotropic with an association with minimum SpO_2_ during sleep (Figure 3B). The lead SNP for *MMP15* (rs4784886) is marginally associated with minimum SpO_2_ in the validation cohort (nominal *p* = 0.051). Overall, of the eight identified genes, four genes (*MMP15*, *SESN1*, *FOXO3* and *BRD3*) are known to be induced by hypoxia and oxidative stress, and three genes (*FOXO3*, *BRD3* and *GSDMA*) are implicated in inflammatory responses. The colocalization of associations to gene expression in specific cell types, clinical and sleep apnea traits suggest biological hypotheses of the pathophysiology underlying OSA.

**Table 2.** Candidate causal chains. Each row represents a SNP with links across an OSA trait, clinical trait and gene expression trait. The link between clinical and OSA traits were thresholded at FDR < 0.2, and the link between clinical and expression traits were thresholded at FDR < 0.05. AHI stands for Apnea-Hypopnea Index. Coordinates correspond to hg19. The eQTL p-value refers to the association p-value of the SNP to the gene expression trait of the gene in the tissue/cell type indicated. The JLIM p-value corresponds to the test for the shared effect between the OSA and clinical traits (center columns) or between gene expression and the clinical trait (right columns).

| SNP | Coordinate | Clinical trait | OSA trait | JLIM<br>p-value | JLIM<br>FDR | Tissue /<br>Cell type | Gene | eQTL<br>p-value | JLIM<br>p-value | JLIM<br>FDR |
| --- | --- | --- | --- | --- | --- | --- | --- | --- | --- | --- |
| rs11153147 | 6:109,304,058 | Mean corpuscular volume | Avg O2<br>saturation | 1.1E-02 | 0.17 | T-cell | *SESN1* (Sestrin 1) | 1.4E-35 | 5.0E-05 | 0.01 |
|  |  |  |  |  |  | Muscle | *SESN1* (Sestrin 1) | 2.6E-05 | 5.0E-05 | 0.05 |
|  |  |  |  |  |  | T-cell | *FOXO3* (Forkhead box O3) | 1.7E-11 | 5.0E-05 | 0.01 |
| rs4784886 | 16:58,065,459 | FEV1/FVC | Min O2<br>saturation | 3.4E-04 | 0.03 | Lung | *MMP15*<br>(Matrix metalloproteinase 15) | 3.3E-10 | 5.0E-05 | 0.02 |
| rs10160596 | 11:65,351,364 | Mean corpuscular<br>hemoglobin concentration | Event Duration | 1.0E-02 | 0.17 | Lung | *SCYL1*<br>(SCY1 like pseudokinase 1) | 5.1E-04 | 5.0E-05 | 0.02 |
| rs146671954 | 9:136,934,203 | Red cell distribution width /<br>Sum eosinophil basophil<br>counts | AHI | 6.9E-04 | 0.02 | Lung | *BRD3*<br>(Bromodomain containing 3) | 2.2E-07 | 1.0E-04 | 0.02 |
| rs2277339 | 12:57,146,069 | Plateletcrit | AHI | 1.0E-02 | 0.15 | monocyte | PRIM1 (DNA primase subunit 1) | 3.2E-06 | 5.0E-05 | 0.04 |
|  |  |  |  |  |  | T-cell | PRIM1 (DNA primase subunit 1) | 1.5E-05 | 5.0E-05 | 0.01 |
| rs7162943 | 15:89,615,275 | Mean platelet volume | Avg O2<br>saturation | 1.0E-03 | 0.07 | monocyte | *ABHD2* (Abhydrolase domain<br>containing 2, acylglycerol<br>lipase) | 2.9E-15 | 1.0E-04 | 0.04 |
| rs34233420 | 17:38,004,929 | Lymphocyte count | AHI/<br>Min O2<br>saturation | 3.4E-04 | 0.02 | T-cell | *GSDMA*<br>(Gasdermin A) | 2.5E-13 | 5.0E-05 | 0.01 |

**Figure 3.**
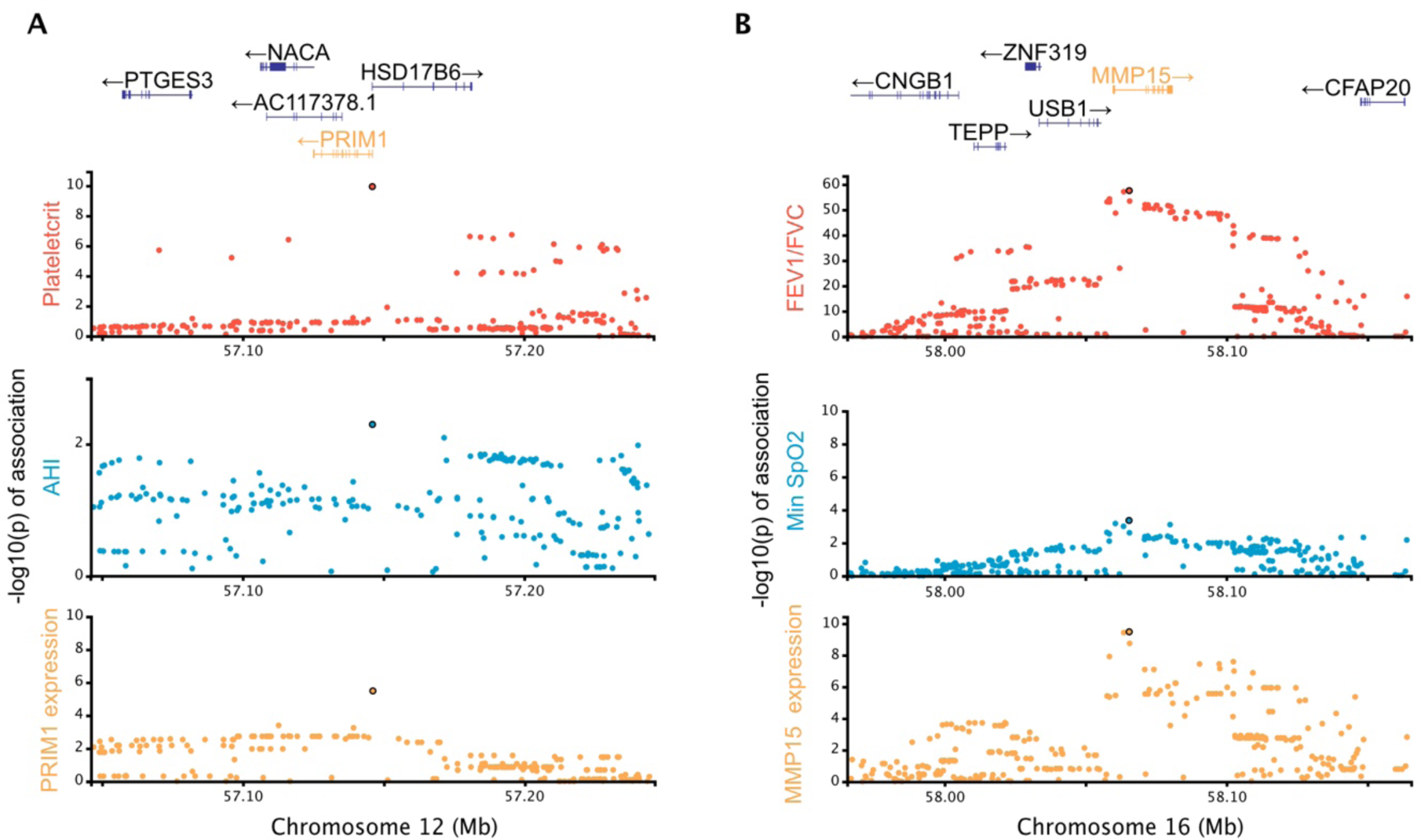
Candidate causal chains linking a clinical trait, OSA trait and gene expression. **A)** A candidate association in chromosome 12 with putative pleiotropic associations between the plateletcrit (clinical trait; red), AHI (OSA trait; blue) and expression of *PRIM1* (DNA primase subunit 1) in monocytes (yellow). **B)** A candidate association in chromosome 16 with putative pleiotropic associations between the clinical trait FEV_1_/FVC (red), minimum O_2_ saturation (blue) and expression of *MMP15* (matrix metallopeptidase 15) in lung (yellow).

We also investigated pleiotropy between gene expression traits and OSA in the three loci harboring known genome-wide significant OSA associations in the discovery sample (Tables 3 and S7). In each locus, we compared OSA trait summary statistics to eQTLs for genes within 1Mb from the most associated variant, where there exists a SNP with eQTL association *p*-value < 5×10^-8^ in the locus. We replicated a previously found pleiotropic effect in a locus on chromosome 17 [24], where minimum oxygen saturation (SpO_2_) colocalizes with expression of the epsilon subunit of the nicotinic receptor (*CHRNE*), in various tissues, including neutrophils, monocytes, spleen and muscle (Figure 4).

**Figure 4.**
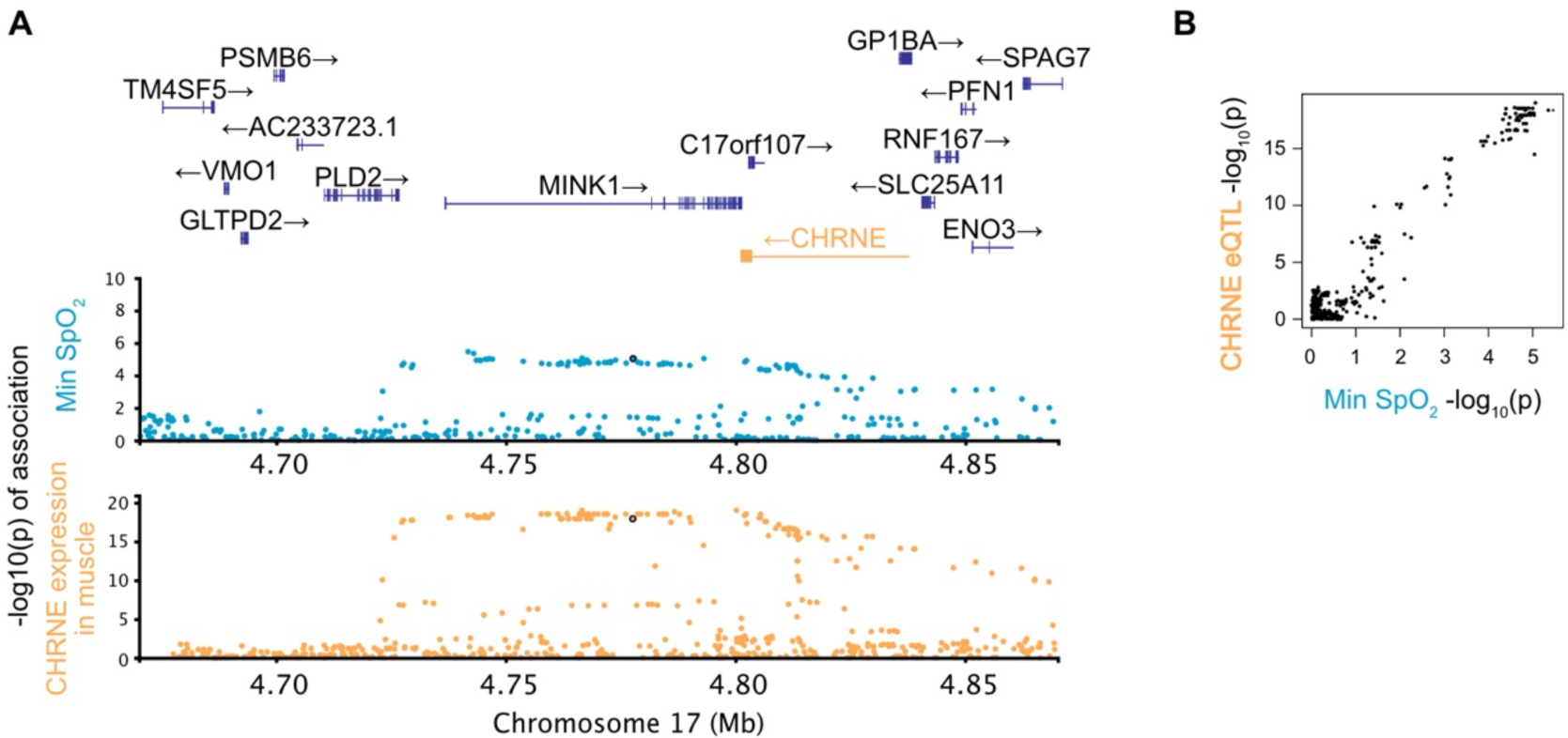
Pleiotropic locus linking gene expression and an OSA related trait. A locus in chromosome 17 has associations between minimum oxygen saturation (blue) and expression of *CHRNE* (cholinergic receptor nicotinic epsilon subunit) in muscle tissues (yellow). Gene expression trait p-values and the appropriate gene in the locus are shown in yellow. Pairwise comparisons of –log_10_(p-values) between associated traits are shown in Panel **B** with matching colors in axis labels.

## Discussion

In our comparison of clinical (respiratory, cardiometabolic, hematologic, inflammatory) traits to OSA-related traits, the strongest finding lies in an intronic region of hexokinase 1 (*HK1*) and is associated with average overnight oxygen saturation level (SpO_2_). This locus is pleiotropic with most of the red blood cell related traits tested (Figure 2) and corresponds to one of the most significant genome-wide associations we had previously reported from this data [24]. Prior to this analysis, two alternative hypotheses for the etiology of this signal had been proposed: that *HK1* acted by modulating inflammation, or that it affected OSA by altering erythrocyte function. Our results provide evidence that is consistent with the erythrocyte pathway hypothesis.

Mutations in *HK1* have been implicated in anemia, together with severe hemolysis and marked decreases in red blood cells [67]. As discussed previously [24], it is possible that *HK1* affects the Rapoport-Luebering shunt through glycolytic pathway intermediates, which in turn mediates oxygen carrying in mature erythrocytes. Factors that influence arterial oxygen levels can lead to a more severe OSA phenotype (i.e., lower average levels of oxygen saturation predispose to greater hypoxemia with each breathing obstruction). Lowered oxygen carrying capacity and thus more tissue hypoxia could also contribute to breathing instability (and thus apneas) via Hypoxia-Inducible Factor-1 (HIF-1) and enhanced carotid body sensitivity and chemoreflex activation, or through long-term respiratory facilitation and plasticity [68,69].

The analysis of pleiotropy can be used to concatenate more than one phenotype to create candidate “causal chains,” which by linking eQTLs to well-powered traits to sleep apnea related traits can hint at promising biological targets. Among the most significant results for this multicomponent model is DNA primase subunit 1 (*PRIM1*). We found that a missense variant in *PRIM1* colocalizes with its own expression levels in monocytes and T cells, plateletcrit, and AHI.

The association of this variant with AHI was replicated in an independent validation cohort. The *PRIM1* deficiency is known to cause lymphopenia along with severe growth retardation, microcephaly, and “triangular face” [70]. The impact of *PRIM1* on craniofacial morphology may contribute to OSA by causing a narrow airway, and thereby increasing the risk of airway obstruction during sleep; however, an inflammatory mechanism is also possible.

Another example of causal chains is matrix metallopeptidase 15 (*MMP15*), a gene whose expression in lung tissue is affected by an eQTL that colocalizes with a lung function phenotype (FEV_1_/FVC) which itself colocalizes with minimum SpO_2_ during sleep. Matrix metallopeptidases (MMPs), a family of proteolytic enzymes that can be activated by inflammation and oxidative stress, participate in and potentiate tissue remodeling by breaking down the extracellular matrix. MMPs have been suggested to play an etiological role in OSA-related pathophysiological responses that may lead to multiple organ dysfunction [71], and specifically, may lead to OSA by contributing to abnormalities in the extracellular matrix of the skeletal muscle in the upper airway, predisposing to passive airway collapse [72]. In particular, *MMP15* (membrane-type 2 MMP) is known to be upregulated by HIF-1α under hypoxic conditions [73,74] and is expressed in alveolar epithelial cells in Idiopathic Pulmonary Fibrosis (IPF) [75], an Interstitial Lung Disease (ILD) characterized by chronic inflammation, progressive formation of scar tissue and decreased lung function. OSA is highly prevalent in ILD as well as associated with subclinical markers of ILD [47]. OSA is also common in Chronic Obstructive Pulmonary Disease (COPD), and this overlap is especially associated with more severe hypoxemia [76]. Our results suggest a common causal pathway linking these lung diseases with OSA.

We also found that *GSDMA* (Gasdermin A) eQTL in T cells colocalizes with associations to the lymphocyte count, AHI as well as oxygen saturation. While Gasdermin A is considered to be expressed mostly in epithelial cells rather than in T cells, broadly, gasdermins mediate inflammatory responses via pyroptosis, a form of programmed cell death leading to the release of proinflammatory molecules, and play a key role in NLRP-inflammasome responses. Increased level of gasdermin D has been shown to mediate hypoxia-related muscle injury in an animal model of OSA [77]. Inflammatory mechanisms may increase risk for OSA through effects on soft tissue structures in the upper airway, muscle function, and/or neural control of breathing, as suggested by a prospective study of inflammation leading to higher risk of OSA [66].

Another interesting result is an eQTL in the epsilon subunit of the nicotinic acetylcholine receptor *CHRNE* that colocalizes with a genome-wide significant association in minimum oxygen saturation. This receptor is present at neuromuscular junctions and mutations in this subunit are known to cause congenital myasthenic syndrome in humans that can result in progressive respiratory impairment [78]. As discussed above, abnormal skeletal muscle responses in the upper airway are considered to be central in the pathogenesis of OSA.

We have shown that we can leverage pleiotropy between OSA and physiological traits to identify the known OSA association in the *HK1* locus. We could not identify two other known OSA associations at 2q12 and 17p11 using this approach, as there is no apparent pleiotropy between OSA and our library of clinical traits. Adding gene expression traits to this analysis increases our discovery ability, as we can identify the 17p11 association as pleiotropic with an eQTL for *CHRNE* in neutrophils, monocytes, spleen and muscle (Table 3). Furthermore, in simulations, we showed that using a large array of clinical trait-associated loci has a potential to boost the power to identify associations in underpowered studies by casting a wider net (Figure 1E). These results warrant further investigation on using gene expressions in diverse cell types and conditions as a library of well-powered traits for underpowered association studies.

**Table 3.**
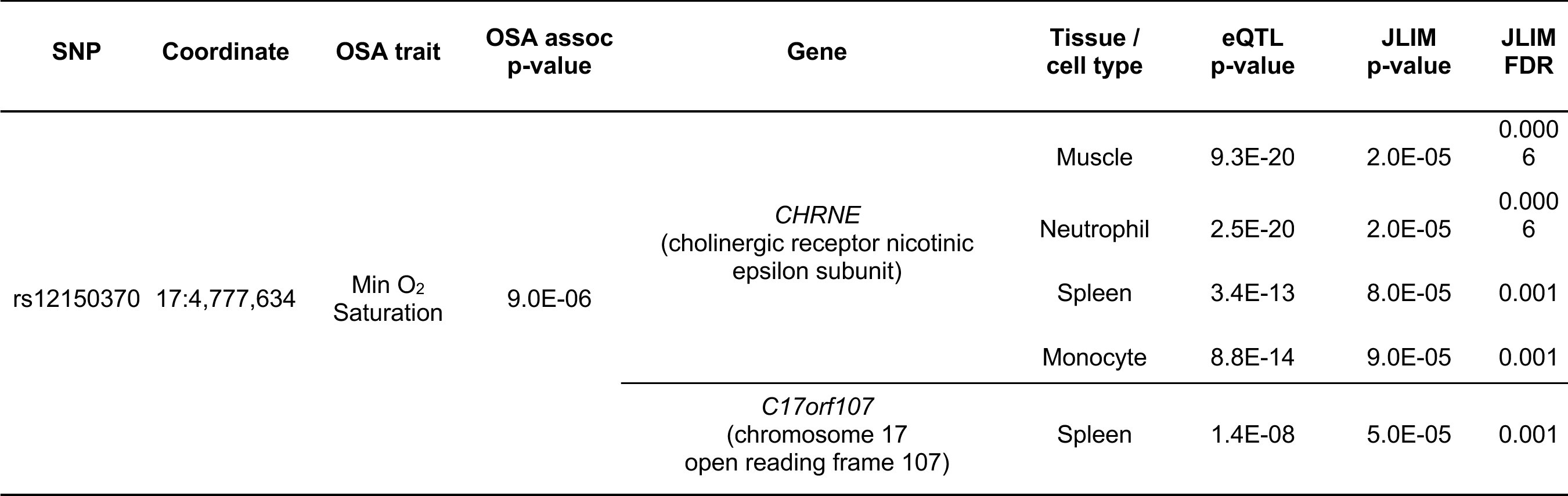
Genome-wide significant loci in OSA traits colocalizing with eQTL. Three loci with known genome-wide significant association to OSA traits were tested for the shared effect with gene expression levels in seven tissue/cell types. The FDR cutoff of 0.05 was applied. The eQTL p-value refers to the association p-value of the SNP to the gene expression trait of the gene indicated, measured in the tissue/cell type indicated. The JLIM p-value refers to a pleiotropy test between the OSA and gene expression traits. See Table S7 for the full list of comparisons tested.

Replication in an independent cohort is frequently necessary to validate GWAS findings in small discovery samples. In this study, we significantly replicate the association of two loci to OSA after Bonferroni correction and identify additional eight loci that replicate only nominally but are probably enriched with true positive associations to OSA (Figure S16). We randomly selected 28,558 unlinked autosomal SNPs (r^2^ < 0.1 within 100kb windows) from our initial OSA GWAS in individuals of European ancestry. We focused on AHI since we detected the most SNPs reaching nominal replication in this trait. We found that 11,910 of these would have been selected for our colocalization analysis, in line with expectation for the presence of a SNP within the 100kb distance satisfying our in-sample threshold of association *p* < 0.01. Out of these, 6,123 had a significant out-of-sample nominal replication (association *p* < 0.05). In comparison, our pleiotropy analysis results for AHI are 4.6-fold enriched at this threshold (4/20 independent hits with r^2^ < 0.1), suggesting the presence of true associations to sleep traits (one-sided Fisher Exact Test *p* = 0.018). In fact, the 76/975 associations that did not show significant evidence of pleiotropy were also slightly enriched for nominal replication relative to the set of randomly selected variants (one-sided Fisher Exact Test *p* = 0.00048, 1.6-fold enrichment). This suggests that additional pleiotropic effects – and therefore true OSA associations – remain to be discovered above the FDR cutoff we applied to our colocalization results although we cannot rule out the possibility that the random SNPs may not fully recapitulate the functional properties of GWAS SNPs from clinical traits.

From a methodological perspective, the analysis of pleiotropy has become an important tool in the analysis of complex trait genetics. Most complex traits are highly polygenic, implying that many variants associated with a single trait will also be associated with other traits or will be in LD with such variants. Different computational methods are required for different applications and for different genetic architectures. If the goal is to increase power to detect an association and the genetic correlation is broadly dispersed over many loci, methods explicitly capitalizing on the broad genetic correlation are capable of producing large power gains [6,29]. In cases where the majority of overlaps in GWAS association peaks between traits are driven by LD between distinct causative variants, power can still be increased with the help of other methods that leverage pleiotropy to reduce multiple testing burden [32,33]. Development of another group of approaches was motivated by the need to link genetic associations to genes via eQTL data [36,38,39] but, as shown here, these methods can be easily adapted to the analysis of other traits. Because of the abundance of association signals, especially for cellular and molecular traits, distinguishing between true pleiotropy due to the same underlying causative variants and different causative variants in LD is important for all the applications. Therefore, in our study of OSA, we selected a method that explicitly models LD structure in the locus. The drawback of this choice is the need to restrict the discovery sample to a demographically homogeneous subset while using the available multi-ethnic cohort for replication. While colocalization tests can distinguish the same and distinct causative variants in overall, the specificity is diminished in cases where association signals of two traits are driven by distinct causative variants in high LD (Figure S2). For example, when the distinct causative variants are in LD between 0.8 and 0.95 (3.7% of simulated data assuming the random LD distribution), eCAVIAR and JLIM misclassify 1.8 and 2.3% of H_2_ loci, respectively, as driven by the same variant (JLIM *p* < 0.01 or at equivalent posterior probability threshold for eCAVIAR). JLIM estimates accurately calibrated *p*-values under H_0_ and conservative approximate *p*-values under H_2_. For underpowered GWAS studies, with limited sample sizes, we lowered JLIM’s genetic resolution parameter *θ* to 0.5 in this study. JLIM does not attempt to distinguish distinct causative variants beyond its specified genetic resolution from the same causal effect. When distinct causative variants for two traits are separated with LD *r^2^* < 0.5, JLIM maintains the ability to distinguish the same and distinct effects in simulations (Figure S2 D-F). And the overall specificity of JLIM to distinguish H_1_ from H_2_ is similar or higher compared to other colocalization method such as eCAVIAR in our simulated datasets (Figure S2B).

Pleiotropy does not necessarily imply a causal relationship between phenotypes. Confounding by unadjusted covariates in GWAS data can further complicate the interpretation of causal chains due to the difficulty to distinguish between direct and indirect associations. Nonetheless, as we demonstrate here, a shared genetic basis between OSA and organismal, cellular and molecular traits can reveal new aspects of the underlying biology. This will likely be of benefit for other clinically relevant traits that are difficult to study at the scale required for GWAS. Traits that are burdensome or expensive to phenotype, rare diseases that are hard to sample and diseases that affect under-represented populations could all lead to underpowered genetic studies, which are unlikely to get dramatically higher sample sizes in the near future. Therefore, there is an unmet need to optimize the signals that can be extracted from small GWAS and the strategy presented here should help in achieving this goal.

## Materials and Methods

### OSA Cohorts

To study pleiotropic associations underlying the risk of OSA, we prepared two sets of cohorts: the discovery cohorts to identify pleiotropic variants and independent replication cohorts to validate their associations to OSA traits. For the discovery cohorts, we used individual-level genotype data in order to determine the significance of pleiotropy by permutation (JLIM) [39]. At the replication stage, we do not carry out any pleiotropy analysis, we only check for genetic association to OSA, so summary-level association statistics were sufficient. In addition, we restricted the genetic ancestry of GWAS discovery cohorts to that of European ancestry, to match GWAS of clinical traits. This was in order to avoid potential issues due to the mismatch of LD patterns in our pleiotropy analysis. In contrast, we did not require the replication cohorts to have any specific ancestry. Thus, our replication cohorts included all available ethnicities.

The discovery cohorts included the subset of samples of European ancestry from the following five cohorts: the Atherosclerosis Risk in Communities Study (ARIC) [79], Osteoporotic Fractures in Men (MrOS) Study [80], Multi-Ethnic Study of Atherosclerosis (MESA) [81], Cardiovascular Health Study (CHS) [82], and the Western Australian Sleep Health Study (WASHS) [83]. ARIC is a study that investigates atherosclerosis and cardiovascular risk factors. It is one of the cohorts included in the Sleep Heart Health Study, which collected polysomnography and genotype data [84]. Genotype data were obtained through dbGaP (phg000035.v1.p1). MESA is a population-based study focused on cardiovascular risk factors, which included participants of four ethnicities: African-, Asian-, European- and Hispanic/Latino-Americans ranging from ages of 45 to 86 years old. We only included samples from European-Americans in the discovery cohort. Polysomnography data measuring sleep-related traits was later obtained from individuals who did not use overnight oxygen, CPAP or an oral device for sleep apnea [85].

MrOS is a multi-center prospective epidemiological cohort assembled to examine osteoporosis, fractures and prostate cancer in older males [86]. An ancillary study (MrOS Sleep) measured sleep disturbances and related outcomes [87]. CHS is a cohort aimed to study coronary heart disease and stroke in individuals aged 65 and older, and genotype data were obtained through dbGaP (Illumina CNV370 and IBC; phg000135.v1.p1 and phg000077.v1.p1). WASHS is a clinic-based study designed to examine OSA and its associated genetic risk factors in patients referred to a sleep clinic in Western Australia. Not all individuals had measurements for the four OSA-related traits of interest. Details on genotyping, imputation and QC procedures have been previously reported [24]. See Table S1 for details.

The replication cohorts were: the Hispanic Community Health Study/Study of Latinos (HCHS/SOL) [88,89], Starr County Health Studies (Starr) [90], Cleveland Family Study (CFS) [91] and Framingham Heart Study (FHS) [92], in addition to non-European samples of CHS and MESA. HCHS/SOL is a population-based study to examine protective and risk factors for many health conditions among Hispanic/Latinos living in four urban areas within the USA (Chicago IL, Miami FL, San Diego CA and Bronx NY). Starr is a cohort collected to study risk factors for diabetes in a population of Mexican-Americans in Texas, later phenotyped for sleep traits [93].

CFS is a family-based study, which recruited patients with OSA, their relatives, and neighborhood control families to study the familial and genetic basis of sleep apnea (356 families of African American or European American ancestry). We included only unrelated individuals from CFS. FHS is an epidemiological cohort established to study cardiovascular disease risk factors, using follow-up medical examinations every two years for the population of European Ancestry in Framingham, MA. Data from the first Sleep Heart Health Study was obtained between 1994-1998. Genotype data were obtained through dbGaP (Affymetrix 500k; phg000006.v7). See Table S6 for the details of each cohort.

We examined the following four OSA-related traits in the discovery and replication cohorts: minimum and average oxygen saturation (SpO_2_), apnea-hypopnea index (AHI) and event duration. Briefly, the minimum and average SpO_2_ were calculated from oximetry-based SpO_2_ measurements over the entire recorded sleep interval excluding occasional waking periods. AHI was scored by counting the number of episodes of complete (apnea) or partial (hypopneas) airflow reduction associated with ≥ 3% desaturation per hour of sleep. The event duration was measured for the average length of apneas and hypopneas, from the nadir of the first reduced breath to the nadir following the last reduced breath (in seconds). The full description of phenotyping protocols are present in the original studies which first reported their genetic analysis in the context of OSA [24–26]. We rank-normalized all OSA traits, separately in each cohort, in order to obtain normally distributed phenotypes.

### Clinical Trait Data

For clinical traits, we used GWAS summary statistics calculated for various traits in the UK Biobank [56,94], blood cell-related phenotypes in a general UK population [55], and cardio-metabolic phenotypes in individuals of European ancestry [58]. There is no sample overlap between clinical trait GWAS data and our discovery or replication cohorts. The full list of clinical traits is shown in Table S2. The GWAS summary statistics for UK Biobank traits and blood cell counts were downloaded from their websites. The summary statistics of cardio-metabolic traits from [58] were obtained directly from the authors.

This research was approved by Partners Healthcare IRB (protocol #2010P001765).

### Identifying pleiotropic variants affecting both clinical traits and OSA

We applied Joint Likelihood Mapping (JLIM version 2.0) [39] to test whether the association signals of clinical and OSA traits were driven by a shared genetic effect. We ran JLIM only on the loci in which there was strong evidence of association to a clinical trait (genome-wide significant) and a suggestive association to the OSA trait (*p* < 0.01 at any SNP in the locus). In these loci, JLIM compares the likelihood of observed association signals under the following three competing possibilities: the OSA trait has no causative variant in the locus (“H_0_”), the same OSA causative variant is shared with the clinical trait (“H_1_”), and the OSA causative variant is distinct from the clinical trait causative variant (“H_2_”), as shown in Figure S1. Since underlying association data have the limited genetic resolution, we test for modified hypotheses *H*_1_^θ^ and *H*_2_^θ^ instead of H_1_ and H_2_. *H*_1_^θ^ represents that the causative variants of two traits are identical or in high LD (*r*^2^ > θ), and *H*_2_^θ^ assumes that the causative variants are separated by the LD threshold (*r*^2^ < θ). In our simulation of underpowered GWAS (see below for the details), only a small fraction (9.0%) of loci had lead SNPs in tight LD (*r*^2^ > 0.8) with causative SNPs; only 15.7% of loci had the modest LD of *r*^2^ > 0.5 between lead and causative SNPs. This contrasts with well-powered GWAS data, for which, in 91.4% of simulated loci, lead SNPs were in LD of *r*^2^ > 0.8 with causative SNPs. To account for this limited statistical resolution of underpowered GWAS data, we lowered *θ* to 0.5 from the default of 0.8 in this study.

JLIM calculates the ratio between the likelihood of the data under *H*_1_^θ^ compared to that under *H*_2_^θ^ and evaluates the significance of this statistic by permuting the phenotypes simulating the lack of causal effect under H_0_. The false positives due to *H*_21_^θ^ are indirectly controlled by the asymptotically conservative behavior of JLIM: with sufficiently large effect sizes or sample sizes, the cumulative distribution function of JLIM statistic shifts lower under *H*_2_^θ^ than under *H*_0_ as previously shown analytically [39]. Thus, under the asymptotic condition, JLIM guarantees that the p-value estimated under *H*_0_ can be used to control for *H*_1_^θ^ as well. However, under non-asymptotic conditions, in particular in case of underpowered cohorts of limited sample sizes, the power to distinguish *H*_1_^θ^ from *H*_2_^θ^ will be diminished. This reduced specificity is more pronounced when the LD between distinct causative variants for two traits is substantial (See Supplements of [39] for the detail). JLIM assumes that only up to one causative variant is present for each trait in a locus. However, simulations showed that the accuracy of JLIM remained robust in the presence of multiple causative variants in a locus (Figure S12).

To run JLIM, we used the genetic association statistics of OSA traits calculated over all common SNPs in a 200kb analysis window around the focal SNP (the lead SNP of a clinical trait). We derived these statistics from our discovery cohorts by combining association signals of each cohort using an inverse-variance weighted meta-analysis approach. The association statistics were calculated in individual cohorts by linear regression adjusting for age, sex, BMI and the top three principal components. The principal components were calculated from genome-wide genotype data in each cohort separately. We used mean imputation for missing covariate values. Multi-allelic SNPs and rare variants with minor allele frequencies (MAF) below 0.05 were excluded from the analysis. We only used variants present in all of the discovery cohorts. To reflect the limited resolution for fine-mapping of causative variants in underpowered studies, we relaxed JLIM’s genetic resolution parameter *θ* to 0.5 from the default value of 0.8. All other parameters were set to the default. We ran JLIM only in the discovery cohorts.

JLIM 2.0 requires individual-level genetic data for underpowered traits in order to run permutations. This limits the applicability of our approach to more general cases where only summary-level data are available. The direct permutation on individual-level genetic data, however, enables robust estimation of p-values even under the vagaries of imputation noise across cohorts for underpowered traits. For the permutation procedure, we used the same pipeline described above to generate permuted association statistics for JLIM. OSA phenotypes were randomly shuffled in each cohort separately. For each permutation, the association statistics were calculated in the same way including all the covariates. Then, the cohort-level association statistics calculated on permuted data were combined across cohorts by meta-analysis. This permutation procedure was repeated up to 100,000 times, adaptively, to estimate JLIM p-values.

We accounted for the multiple testing burden with the False Discovery Rate (FDR), separately for each clinical and OSA trait combination. Specifically, we used the Benjamini-Hochberg procedure to calculate the FDR as follows:

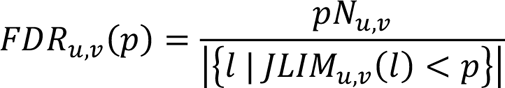

where *u* and *v* indicate a combination of clinical and OSA traits, *JLIM_u,v_*(*l*) is the JLIM p-value of tested locus *l*, and *_u,v_* is the number of loci tested between the trait pairs *u* and *v*. JLIM hits were obtained at the FDR threshold of 0.2. The JLIM/cFDR consensus was defined by the intersection between the JLIM (FDR < 0.2) and cFDR hits defined at the same p-value threshold. Specifically, given the JLIM p-value cutoff of FDR 0.2, i.e., *p*_0.2_ such that *FDR_u,v_*(*p*_0.2_) = 0.2, the cFDR hits were defined by the association p-value below *p*_0.2_ for an OSA trait. Note that the p-value of association to a clinical trait is always ascertained to be less than 5 × 10^-8^ in our analysis. The cFDR association p-values were examined at the focal SNP.

### Comparison with eCAVIAR

eCAVIAR (version 2.2) was run in the default setting. The reference LD matrix was set to the LD estimated from the subjects of European ancestry (n=10,000) randomly subsampled out of our OSA discovery cohort. Colocalization Posterior Probability (CLPP) was calculated for lead SNPs of well-powered clinical traits to identify pleiotropic loci. The CLPP thresholds were calibrated to be equivalent to P-value cutoffs using unfiltered null simulation datasets (H_0_), which were not preconditioned on the minimum association P-values in loci.

### Replication of OSA associations in independent samples

We validated the OSA associations identified in the discovery cohorts by replicating them in out-of-sample multi-ethnic replication cohorts (Table S6). There was no sample overlap between our discovery and replication cohorts. We combined the *p*-values of associations across the six replication cohorts by applying an inverse variance-weighted meta-analysis technique. We defined the *p*-value of association < 0.05 as nominal evidence of replication and the *p*-value < 0.05/61 as a more stringent Bonferroni-corrected replication cutoff, given that 61 independent SNPs were uncovered for their pleiotropic associations to OSA in the discovery cohort (Table 1, Table S5).

### Comparison of nominal replication rates with random SNPs

To assess the sensitivity of our pleiotropy analysis, we compared the nominal replication rate of random SNPs with that of candidate pleiotropic variants identified in the discovery cohort. We started with 87,938 SNPs randomly selected across the autosomes, excluding chromosome 6 to avoid the major histocompatibility complex locus. Then, we used Plink to apply LD pruning on the random set of SNPs and obtained a subset of 28,558 independent SNPs. The LD pruning procedure ensured that the R^2^ between SNPs was less than 0.1 in the distance of 100kb in the LD background of non-Finnish Europeans (n=404) from the 1000 Genomes Project. For the fair comparison between random and predicted pleiotropic SNPs, we further filtered these random SNPs based on the OSA association p-values in 200kb windows. Overall, we kept only 11,910 random independent SNPs that have an AHI-associated SNP (P < 0.01) within 100kb distance. On all loci examined for the pleiotropy, we similarly applied the LD pruning procedure on the focal SNPs to obtain a subset of independent SNPs. During the pruning steps, SNPs with more significant JLIM p-values were preferentially retained. After the LD pruning, we obtained 995, 1,024, 967 and 961 independent SNPs for AHI, average SpO_2_, minimum SpO_2_, and event duration, respectively; of which 20, 12, 6 and 11 SNPs are predicted to be pleiotropic in the discovery cohort.

### Identifying pleiotropic variants affecting both gene expressions and OSA

We used *cis*-expression quantitative trait loci (eQTLs) from the Gene-Tissue Expression project (GTEx release v8) [63] and BLUEPRINT epigenome project [64], to examine pleiotropy between the variation in gene expression levels and OSA phenotypes. Among the GTEx datasets, we only considered liver (n=178), spleen (n=179), skeletal muscle (n=588) and lung (n=436) tissues for our analysis, based on the potential relevance of these tissues to OSA and their sample sizes. Again, we used eQTLs calculated only with samples of European ancestry for this analysis. The genome-wide summary statistics of European American eQTLs were obtained from the GTEx Consortium. For the analysis of immune cell eQTLs, we used BLUEPRINT datasets which consisted of genotypes of participants and expression profiles of CD14^+^ monocytes (n=191), neutrophils (n=196) and CD4^+^ T cells (n=167). The RNA transcripts of BLUEPRINT samples were derived from unstimulated primary cells collected from healthy individuals of European ancestry. The genome-wide summary statistics of BLUEPRINT eQTLs were downloaded from the eQTL Catalogue [95].

We start with the 61 index SNPs corresponding to the candidate pleiotropic loci identified by JLIM FDR < 0.2 and cFDR consensus (Table S5). Using GTEx and BLUEPRINT eQTLs, we scanned for pleiotropy between eQTLs and clinical traits. We considered all protein-coding genes whose transcription start sites (TSS) were less than 1Mb away from the focal SNP of a clinical trait (1,011 unique genes; 5,690 eQTLs). The protein-coding genes were defined by Ensembl annotation (release 104). The genes with eQTL association *p*-value > 0.05 at all SNPs in the locus were excluded due to weak evidence of association to gene expression. Overall, a total of 5,860 clinical traits/eQTLs pairs (1,009 unique genes; 5,023 eQTLs) were tested for the pleiotropy using JLIM in the default setting (Table 2). JLIM version 2.5 was used to test for pleiotropy only using summary-level eQTL data that are publicly available. The p-values were calculated by adaptive resampling (up to 10,000 iterations). The null distribution of JLIM statistic was generated by random sampling of phenotypes from a normal distribution, sampling of genotypes from the reference LD panel (1000 Genomes, non-Finnish Europeans), and then linearly regressing the sampled phenotypes on the genotypes. Multiallelic SNPs and rare variants with MAF < 0.05 were excluded from the analysis. The multiple testing burden was accounted for with the False Discovery Rate (FDR) by applying the Benjamini-Hochberg procedure separately to each tissue/cell type and clinical trait combination.

### Using gene expression as clinical traits

In three loci with known genome-wide associations to OSA traits, we tested for pleiotropy between gene expression and OSA traits using *cis*-eQTLs as clinical traits. We examined four GTEx tissues and three BLUEPRINT immune cell types, similarly to the above causal chain analysis. There are a total 57 eQTLs in the three loci satisfying: 1) the eQTL p-values < 5 × 10^-8^ for a SNP within the 200kb window centered at the known OSA-associated SNP, and 2) the transcription start sites being within 1 Mb from the OSA-associated SNP. Using these 57 eQTLs as clinical traits, we applied JLIM in the default setting (Table S7). For the OSA traits, we used the association statistics from our discovery cohorts. The JLIM FDR was calculated using the Benjamini-Hochberg procedure (a total of 57 tests).

### Simulated datasets

To compare the accuracy of JLIM to other methods, we simulated genetic loci with pleiotropic associations under different scenarios with unbalanced sample sizes. For one of two traits, we simulated a well-powered GWAS of a quantitative trait with a sample size of 150,000. For the other trait, we simulated an under-powered GWAS with a much smaller sample size of 10,000. To generate datasets of realistic LD backgrounds, we used real genotypes of 80 randomly picked loci across the genome. Each locus was 200kb in length. Chromosome 6 and the sex chromosomes were excluded from our simulations due to the difference in LD patterns from the rest of genome.

In these loci, we generated 80,000 sets of association statistics under each of the following six scenarios (H_0_, H_1_ and four configurations of H_2_). For the simulations of H_0_, the SNP in the midpoint of the genomic segment was chosen as the causative variant for well-powered traits, but no causal effect was simulated for underpowered traits. For H_1_, the midpoint SNP was taken as the shared causative variant for both well-powered and underpowered traits. For H_2_, the midpoint SNP was selected to be causal for well-powered traits, but a distinct SNP was selected as the causative variant for underpowered traits. The distinct SNP was randomly chosen in the intervals of LD relative to the midpoint SNP, selecting one in each of the following LD ranges: |r| < 0.3, 0.3 < |r| < 0.6, 0.6 < |r| < 0.8 and 0.8 < |r| < 0.95.

For each pair of causative variants, we randomly sampled the genetic effect sizes (*α*_1_, *β*_2_), corresponding to well-powered and underpowered traits, respectively, from the following bivariate normal distribution:

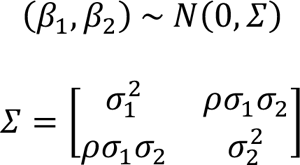

where *σ*_1_^2^ and *σ*_2_^2^ are per-SNP heritability of two traits, set to 5 × 10^-5^ under the assumptions of the heritability of 0.5, causal fraction of 0.01 and 1,000,000 independent markers across the genome. The parameter *ρ* was used to represent the correlation of causal effect sizes between two traits. For H_1_, we generated simulated data under no correlation (*ρ* = 0) as well as moderate to high correlation (*ρ*= 0.5, 0.7 or 0.9). For H_2_, we assumed *ρ* = 0 since the causative variants are not shared. For H_0_, *β*_2_ was set to 0.

JLIM only requires GWAS summary statistics for the well-powered trait. Therefore, for all SNPs *j* = 1, … , *m* in each locus, we generated summary statistics by sampling the observed association statistics *z* = (*z*_1_, …, *z_j_*, …, *z_m_*) from the following multivariate normal distribution [52]:

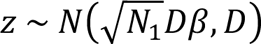

where *N*_1_ is the GWAS sample size (150,000), *m* is the number of markers in the locus, *D* is the *m* × *m* local LD matrix, and *β* is the *m*-dimensional vector of true causal effects (on standardized genotype values) of all SNPs in the locus. Here, *β* was set to *β*_1_ at the causative SNP and to 0 at all other SNPs. The *p*-values of association were calculated from the *z* scores as follows:

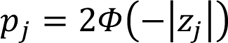

where *z_j_* is the association statistic at SNP *j* and *Φ* is the standard normal cumulative distribution function.

JLIM 2.0 requires individual-level genotype data as well as association statistics for the second trait. Therefore, for the underpowered trait, we generated a discovery cohort with simulated phenotypes for all individuals; then, we calculated the association statistics by linear regression between genotypes and phenotypes, instead of sampling the summary statistics from a multivariate normal distribution. The genotype data of 10,000 individuals were obtained by subsampling from six cohorts of European Ancestry (MESA, ARIC, MrOS, CHS, CFS, FHS and WASHS). The phenotype value *y_i_* of each individual *i* was generated by random sampling from the following standard normal distribution:

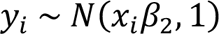

where *x_i_* is the genotype of the causative variant in the individual *i* (standardized to the mean of 0 and the variance of 1). The association statistics and p-values of association were obtained by regressing the phenotypes *y_i_* on the genotypes of each SNP in the locus. We excluded multiallelic sites and variants with MAF < 0.05 from the analysis.

The above process generated up to 480,000 sets of association data under scenarios of H_0_, H_1_ and H_2_ in the genetic background of 80 loci (i.e., 80 × 6 × 1,000). Because of sampling noise, in a minority of simulations, association data of some loci failed to pass the genome-wide significance threshold for well-powered traits. We rejected such simulation runs to mimic our data analysis, where we only included loci in which the clinical trait association p-value was genome-wide significant. Similarly, we excluded instances where there exists no SNP with the p-value association < 0.01 for underpowered traits. In total, we retained 25,958, 31,858, 28,602, 29,131, 27,025, and 25,115 sets of association data for H_0_, H_1_, and H_2_ of distinct causative variants in low to high LD, respectively.

Next, we estimated the distribution of LD between random pairs of SNPs in the 80 simulated loci and titrated the causative variants of H_2_ simulations to follow this distribution. Specifically, we subsampled H_2_ datasets to fit to the following composition: 72% for |r| < 0.3, 19% for 0.3 < |r| < 0.6, 5% for 0.6 < |r| < 0.8 and 4% for 0.8 < |r| < 0.95 (Figure S2C). After this, we generated GWAS datasets of *T* loci by random subsampling of H_0_, H_1_ and H_2_. *T* represents the total number of GWAS loci available from well-powered traits and varied from 500 to 5,000. The proportion of H_0_ out of *T* loci varied from 0 to 30%. The relative ratios of loci simulating H_1_ and H_2_ were set to 1:19, 1:9, 1:4 or 1:0.

To assess the power to replicate the pleiotropic associations in independent validation cohorts, we calculated the expected association statistic *Z_j_^v^* for the underpowered trait at the focal SNP *j*:

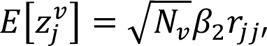

where the focal SNP *j* was defined by the lowest p-value of association to the well-powered trait, *N_v_* is the sample size of validation cohort for the underpowered trait, ranging from 10,000 to 35,000, *β*_2_ is the effect size for the underpowered trait at its causative SNP *j*′, and *r_jj_*, is the LD between two SNPs *j* and *j*′. For the LD backgrounds of validation cohorts, we used CEU and YRI from the 1000 Genomes Project to represent ancestry-matched and trans-ethnic validation cohorts, respectively. The expected number of replicated signals after Bonferroni correction was calculated by estimating:

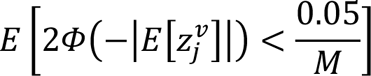

where *M* is the number of pleiotropic associations found in each subsampled GWAS dataset of *T* loci.

Last, we also evaluated the accuracy of our method under the presence of multiple causative variants that were shared between two traits in H_1_. In addition to the midpoint SNP, we randomly picked another SNP in the locus as a second shared causative variant. The effect sizes of the two causative variants for well-powered and underpowered traits were sampled from the same bivariate normal distribution. We generated GWAS datasets for the well-powered and underpowered traits in a similar manner as the simulations of single causative variants. The correlation of effect sizes between well-powered and underpowered traits (*ρ*) was set to 0.7 for H_1_. The proportion of H_0_ was assumed to be 30% of examined GWAS loci (2,500). The relative proportion of H_1_ and H_2_ was set to 1:19. The proportion of loci with two causative variants was set to 1/4 of all H_1_ as expected under Poisson distribution with the causal fraction of 0.01.

### Simulated datasets for the meta-analysis of more than two traits

GWAS data were simulated for ten well-powered and one underpowered traits under H_0_, H_1_ and H_2_ (Figure S15A). One of the well-powered traits was ascertained to have a genome-wide significant association (the main well-powered trait). Here, we simulated association statistics only for the causative SNP of the main well-powered trait rather than simulating for entire SNPs in the locus. For simplicity of simulation, we assumed that the lead SNP of the main well-powered trait is in tight LD with its causative variant. The rest of well-powered traits have the same or distinct causative variants from the main well-powered trait. The number of additional well-powered traits simulating the same causative variants was randomly decided by sampling from a binomial distribution *Binom* 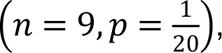 and the rest were assumed to have distinct causative variants. For underpowered traits, the causative variant was assumed to be absent, same or distinct from the main well-powered trait depending on whether the locus was simulated under H_0_, H_1_ or H_2_, respectively. Effect sizes of all traits sharing the same causative variant were sampled together from a multivariate normal distribution with the correlation of effect sizes between traits set to 0.7. Effect sizes of traits simulating distinct causative variants were sampled independently and then multiplied by a random variable representing the LD between causative and tested SNPs. The association statistics were generated with the sample sizes of n=150,000 for well-powered traits and n=10,000 for underpowered traits. In total, 3,751, 613, and 6,229 sets of simulated association statistics were generated under H_0_, H_1_ and H_2_, respectively. From these simulated H_0_, H_1_ and H_2_ datasets, GWAS datasets of 2,500 loci were generated by random subsampling at the proportions of 30%, 3.5% and 66.5%, respectively (H_1_:H_2_ ratio of 1:19). We evaluated the power to replicate the candidate pleiotropic loci using the subsampled GWAS datasets (1,000 iterations). The power to replicate after Bonferroni correction was estimated in the same way as other simulations. The validation cohort were assumed to be of the same genetic ancestry as the discovery cohort, and the sample sizes varied from n=10,000 to 35,000. For pairwise analyses, we used only the main well-powered trait and underpowered trait. On the other hand, for full multi-trait analyses, we used the entire data including all ten well-powered traits and one underpowered trait.

### Meta-analysis methods

We benchmarked the power to replicate pleiotropic associations in underpowered cohorts by applying three meta-analysis methods - MetABF, CPBayes and iGWAS - on simulated datasets. We ran MetABF in the subset-exhaustive mode to scan all possible subsets of pleiotropic traits. The parameter for the correlation of effect sizes was set to the known value of simulation (*ρ* = 0.7). MetABF was run three times with the scale parameter of causal effect sizes set to 0.1, 0.2 and 0.4, and the Approximate Bayes Factors (ABF) were averaged over the three runs as recommended by the authors. MetABF reports the ABF relative to H_0_, thus we calculated the ABF of pleiotropy to the underpowered trait relative to that of no pleiotropy to the underpowered trait by summing ABFs over subsets. In contrast, we ran CPBayes in the fully automatic setting. All parameters of the pleiotropy were directly learned from the data. We used the PPAj (Posterior Probability of Association) of the underpowered trait to identify pleiotropic associations. In comparison to the two Bayesian methods, iGWAS is a non-parametric meta-analysis test. We ran iGWAS in the default setting. For all three methods, candidate pleiotropic associations were identified in the discovery cohort at the same p-value cutoff of 0.01. For the Bayesian methods, we applied the equivalent ABF and PPAj thresholds by calibrating them to have the false positive rate of 0.01 in our null simulation (H_0_).

## Supporting information

Revision Supplementary Figures

Revision Supplementary Tables

## Acknowledgments

This work was supported by the National Institute of Health grant R01-HL113338. Susan Redline is partially supported by grants from the National Institutes of Health [K01-HL135405, R01-HL113338, R35-HL135818]. Shamil Sunyaev is partially supported by the National Institute of General Medical Sciences grant R35-GM127131 and the National Institute of Health grant R01-MH_1_01244.

The Atherosclerosis Risk in Communities (ARIC) study has been funded in whole or in part with Federal funds from the National Heart, Lung, and Blood Institute, National Institutes of Health, Department of Health and Human Services (contract numbers HHSN268201700001I, HHSN268201700002I, HHSN268201700003I, HHSN268201700004I and HHSN268201700005I), R01HL087641, R01HL059367 and R01HL086694; National Human Genome Research Institute contract U01HG004402; and National Institutes of Health contract HHSN268200625226C. The authors thank the staff and participants of the ARIC study for their important contributions. Infrastructure was partly supported by Grant Number UL1RR025005, a component of the National Institutes of Health and NIH Roadmap for Medical Research.

MESA and the MESA SHARe project are conducted and supported by the National Heart, Lung, and Blood Institute (NHLBI) in collaboration with MESA investigators. Support for MESA is provided by contracts HHSN268201500003I, N01-HC-95159, N01-HC-95160, N01-HC-95161, N01-HC-95162, N01-HC-95163, N01-HC-95164, N01-HC-95165, N01-HC-95166, N01-HC-95167, N01-HC-95168, N01-HC-95169, UL1-TR-000040, UL1-TR-001079, UL1-TR-001420. MESA Family is conducted and supported by the National Heart, Lung, and Blood Institute (NHLBI) in collaboration with MESA investigators. Support is provided by grants and contracts R01HL071051, R01HL071205, R01HL071250, R01HL071251, R01HL071258, R01HL071259, and by the National Center for Research Resources, Grant UL1RR033176. The provision of genotyping data was supported in part by the National Center for Advancing Translational Sciences, CTSI grant UL1TR001881, and the National Institute of Diabetes and Digestive and Kidney Disease Diabetes Research Center (DRC) grant DK063491 to the Southern California Diabetes Endocrinology Research Center.

The Osteoporotic Fractures in Men (MrOS) Study is supported by NIH funding. The following institutes provide support: the National Institute on Aging (NIA), the National Institute of Arthritis and Musculoskeletal and Skin Diseases (NIAMS), NCATS, and NIH Roadmap for Medical Research under the following grant numbers: U01 AG027810, U01 AG042124, U01 AG042139, U01 AG042140, U01 AG042143, U01 AG042145, U01 AG042168, U01 AR066160, and UL1 TR000128. The NHLBI provides funding for the MrOS Sleep ancillary study “Outcomes of Sleep Disorders in Older Men” under the following grant numbers: R01 HL071194, R01 HL070848, R01 HL070847, R01 HL070842, R01 HL070841, R01 HL070837, R01 HL070838, and R01 HL070839. The NIAMS provides funding for the MrOS ancillary study ‘Replication of candidate gene associations and bone strength phenotype in MrOS’ under the grant number R01 AR051124. The NIAMS provides funding for the MrOS ancillary study ‘GWAS in MrOS and SOF’ under the grant number RC2 AR058973. The investigators website can be found at https://mrosonline.ucsf.edu/.

This CHS research was supported by NHLBI contracts HHSN268201200036C, HHSN268200800007C, HHSN268201800001C, N01HC55222, N01HC85079, N01HC85080, N01HC85081, N01HC85082, N01HC85083, N01HC85086; and NHLBI grants U01HL080295, R01HL087652, R01HL105756, R01HL103612, R01HL120393, and U01HL130114 with additional contribution from the National Institute of Neurological Disorders and Stroke (NINDS). Additional support was provided through R01AG023629 from the National Institute on Aging (NIA). A full list of principal CHS investigators and institutions can be found at <u>CHS-NHLBI.org</u>.

The provision of genotyping data was supported in part by the National Center for Advancing Translational Sciences, CTSI grant UL1TR001881, and the National Institute of Diabetes and Digestive and Kidney Disease Diabetes Research Center (DRC) grant DK063491 to the Southern California Diabetes Endocrinology Research Center. The content is solely the responsibility of the authors and does not necessarily represent the official views of the National Institutes of Health.

Funding for the Western Australian Sleep Health Study was obtained from the Sir Charles Gairdner and Hollywood Private Hospital Research Foundations, the Western Australian Sleep Disorders Research Institute, and the Centre for Genetic Epidemiology and Biostatistics at the University of Western Australia. Funding for the GWAS genotyping obtained from the Ontario Institute for Cancer Research and a McLaughlin Centre Accelerator Grant from the University of Toronto.

The Hispanic Community Health Study/Study of Latinos was carried out as a collaborative study supported by contracts from the NHLBI to the University of North Carolina (HHSN268201300001I / N01-HC65233), University of Miami (HHSN268201300004I / N01-HC65234), Albert Einstein College of Medicine (HHSN268201300002I / N01-HC65235), University of Illinois at Chicago (HHSN268201300003I), Northwestern University (N01-HC65236), and San Diego State University (HHSN268201300005I / N01-HC65237). The following Institutes/Centers/Offices contribute to the HCHS/SOL through a transfer of funds to the NHLBI: National Institute on Minority Health and Health Disparities, National Institute on Deafness and Other Communication Disorders, National Institute of Dental and Craniofacial Research, National Institute of Diabetes and Digestive and Kidney Diseases, National Institute of Neurological Disorders and Stroke, NIH Institution-Office of Dietary Supplements. The Genetic Analysis Center at Washington University was supported by NHLBI and NIDCR contracts (HHSN268201300005C AM03 and MOD03). The views expressed in this manuscript are those of the authors and do not necessarily represent the views of the National Heart, Lung, and Blood Institute; the National Institutes of Health; or the U.S. Department of Health and Human Services. The authors thank the staff and participants of HCHS/SOL for their important contributions. Investigator’s website - http://www.cscc.unc.edu/hchs/

The Starr County Health Studies is supported in part by grants R01 DK073541, U01 DK085501, R01 AI085014, and R01 HL102830 from the National Institutes of Health, and funds from the University of Texas Health Science Center at Houston. We thank the field staff in Starr County for their careful collection of these data and are especially grateful to the participants who so graciously cooperated and gave of their time.

The Cleveland Family Study has been supported by National Institutes of Health grants [5-R01-HL046380-15, 5-KL2-RR024990-05, R35 HL135818, and HL113338].

The Framingham Heart Study (FHS) has been supported by contracts N01-HC-25195 and HHSN268201500001I and grant R01 HL092577. The Framingham Heart Study thanks the study participants and the multitude of investigators who over its 70 year history continue to contribute so much to further our knowledge of heart, lung, blood and sleep disorders and associated traits.

This study makes use of data generated by the BLUEPRINT Consortium. A full list of the investigators who contributed to the generation of the data is available from www.blueprint-epigenome.eu. Funding for the project was provided by the European Union’s Seventh Framework Programme (FP7/2007-2013) under grant agreement no 282510 BLUEPRINT.

The Genotype-Tissue Expression (GTEx) Project was supported by the Common Fund of the Office of the Director of the National Institutes of Health, and by NCI, NHGRI, NHLBI, NIDA, NIMH, and NINDS.

## Author Contributions

S.R. Sunyaev, C. Cotsapas, S. Chun, S. Akle, S. Redline conceived and designed the experiment. S. Chun, S. Akle, A. Teodosiadis analyzed the data. B.E. Cade, H. Wang, T. Sofer, D.S. Evans, K.L. Stone, S.A. Gharib, S. Mukherjee, L.J. Palmer, D. Hillman, J.I. Rotter, C.L. Hanis, J.A. Stamatoyannopoulos, S. Redline contributed data/analysis tools. S. Chun, S. Akle, C. Cotsapas, S.R. Sunyaev wrote the original draft. Every author reviewed and edited the manuscript.

## Supporting Information

**Figure S1. Schematic of JLIM analysis.**

**Figure S2. Sensitivity for H_1_ and specificity to distinguish H_1_ from H_2_. Figure S3. Winner’s curse.**

**Figure S4. The number of loci that were identified in simulated discovery and validation cohorts, broken down by the configuration of causative variants.**

**Figure S5. The projected number of replicated loci in simulations, varying the relative ratio between H_1_ and H_2_.**

**Figure S6. The projected number of replicated loci in simulations, varying the proportion of H_0_ loci.**

**Figure S7. The projected number of replicated loci in simulations where the cFDR threshold was tightened to identify the same number of candidate loci as the JLIM/cFDR consensus method in a discovery cohort.**

**Figure S8. The projected number of replicated loci in simulations where the cFDR threshold was calibrated to have the same false positive rate for H_0_.**

**Figure S9. The projected number of replicated loci in simulations where the number of association peaks available from well-powered traits varied.**

**Figure S10. The projected number of replicated loci in simulations where the correlation of effect sizes for H_1_ varied.**

**Figure S11. The projected number of replicated loci in simulations of trans-ethnic replication.**

**Figure S12. The projected number of replicated loci in simulations of multiple causative variants for H_1_.**

**Figure S13. Comparison with Bayesian meta-analysis methods with a validation cohort from the same ancestry.**

**Figure S14. Comparison with Bayesian meta-analysis methods with a validation cohort from the different ancestry (YRI).**

**Figure S15. Simulated power of multi-trait meta-analysis compared to pairwise analysis.**

**Figure S16. Pleiotropic loci identified by the intersection of JLIM and cFDR are enriched with SNPs that nominally replicate out of sample.**

**Table S1. Sample sizes of discovery cohorts used for the pleiotropy analysis. Table S2. Clinical traits used for the pleiotropy analysis.**

**Table S3. Mendelian randomization analysis p-values testing effect of exposure on outcome.**

**Table S4. Overall number of total tests, identified pleiotropic loci, and nominally replicated signals.**

**Table S5. List of identified pleiotropic loci and their replication p-values for the corresponding OSA traits (JLIM FDR < 0.2).**

**Table S6. Sample sizes of validation cohorts used for the replication study.**

**Table S7. Full results of pleiotropy tests between eQTLs and OSA traits in three known OSA GWAS loci at the eQTL p-value < 5e-8.**

## Notes

http://genetics.bwh.harvard.edu/wiki/sunyaevlab/jlim2.0

https://github.com/cotsapaslab/jlim/

### Competing Interest Statement

The authors have declared no competing interest.

### Summary of Updates

-

## References

1. Visscher PM, Wray NR, Zhang Q, Sklar P, McCarthy MI, Brown MA, et al. 10 Years of GWAS Discovery: Biology, Function, and Translation. Am J Hum Genetics. 2017;101(1):5–22.

2. Zhang Y, Qi G, Park JH, Chatterjee N. Estimation of complex effect-size distributions using summary-level statistics from genome-wide association studies across 32 complex traits. Nat Genet. 2018;50(9):1318–26.

3. Bulik-Sullivan B, Finucane HK, Anttila V, Gusev A, Day FR, Loh PR, et al. An atlas of genetic correlations across human diseases and traits. Nat Genet. 2015;47(11):1236–41.

4. Pickrell JK, Berisa T, Liu JZ, Ségurel L, Tung JY, Hinds DA. Detection and interpretation of shared genetic influences on 42 human traits. Nat Genet. 2016;48(7):709–17.

5. Shi H, Mancuso N, Spendlove S, Pasaniuc B. Local Genetic Correlation Gives Insights into the Shared Genetic Architecture of Complex Traits. Am J Hum Genetics. 2017;101(5):737–51.

6. Turley P, Team 23andMe Research, Walters RK, Maghzian O, Okbay A, Lee JJ, et al. Multi-trait analysis of genome-wide association summary statistics using MTAG. Nat Genet. 2018;50(2):229–37.

7. Fortney K, Dobriban E, Garagnani P, Pirazzini C, Monti D, Mari D, et al. Genome-Wide Scan Informed by Age-Related Disease Identifies Loci for Exceptional Human Longevity. Plos Genet. 2015;11(12):e1005728.

8. Peppard PE, Hagen EW. The Last 25 Years of Obstructive Sleep Apnea Epidemiology—and the Next 25? Am J Resp Crit Care. 2017;197(3):310–2.

9. Olaithe M, Bucks RS. Executive Dysfunction in OSA Before and After Treatment: A Meta-Analysis. Sleep. 2013;36(9):1297–305.

10. Gami AS, Hodge DO, Herges RM, Olson EJ, Nykodym J, Kara T, et al. Obstructive sleep apnea, obesity, and the risk of incident atrial fibrillation. J Am Coll Cardiol. 2007;49(5):565–71.

11. Gozal D, Ham SA, Mokhlesi B. Sleep Apnea and Cancer: Analysis of a Nationwide Population Sample. Sleep. 2016;39(8):1493–500.

12. Nieto FJ, Peppard PE, Young T, Finn L, Hla KM, Farré R. Sleep-disordered breathing and cancer mortality: results from the Wisconsin Sleep Cohort Study. Am J Resp Crit Care. 2012;186(2):190–4.

13. Kendzerska T, Gershon AS, Hawker G, Leung RS, Tomlinson G. Obstructive Sleep Apnea and Risk of Cardiovascular Events and All-Cause Mortality: A Decade-Long Historical Cohort Study. Plos Med. 2014;11(2):e1001599.

14. Oldenburg O, Wellmann B, Buchholz A, Bitter T, Fox H, Thiem U, et al. Nocturnal hypoxaemia is associated with increased mortality in stable heart failure patients. Eur Heart J. 2016;37(21):1695–703.

15. Nagayoshi M, Punjabi NM, Selvin E, Pankow JS, Shahar E, Iso H, et al. Obstructive sleep apnea and incident type 2 diabetes. Sleep Med. 2016;25:156–61.

16. BaHammam AS, Kendzerska T, Gupta R, Ramasubramanian C, Neubauer DN, Narasimhan M, et al. Comorbid depression in obstructive sleep apnea: an under-recognized association. Sleep Breath. 2016;20(2):447–56.

17. Torres G, Sánchez-de-la-Torre M, Barbé F. Relationship Between OSA and Hypertension. Chest. 2015;148(3):824–32.

18. Somers VK, White DP, Amin R, Abraham WT, Costa F, Culebras A, et al. Sleep Apnea and Cardiovascular Disease An American Heart Association/American College of Cardiology Foundation Scientific Statement From the American Heart Association Council for High Blood Pressure Research Professional Education Committee, Council on Clinical Cardiology, Stroke Council, and Council on Cardiovascular Nursing In Collaboration With the National Heart, Lung, and Blood Institute National Center on Sleep Disorders Research (National Institutes of Health). J Am Coll Cardiol. 2008;52(8):686–717.

19. Redline S, Yenokyan G, Gottlieb DJ, Shahar E, O’Connor GT, Resnick HE, et al. Obstructive Sleep Apnea–Hypopnea and Incident Stroke. Am J Resp Crit Care. 2010;182(2):269–77.

20. Gottlieb DJ, Yenokyan G, Newman AB, O’Connor GT, Punjabi NM, Quan SF, et al. Prospective Study of Obstructive Sleep Apnea and Incident Coronary Heart Disease and Heart Failure. Circulation. 2010;122(4):352–60.

21. Mukherjee S, Saxena R, Palmer LJ. The genetics of obstructive sleep apnoea. Respirology. 2018;23(1):18–27.

22. Palmer LJ, Redline S. Genomic approaches to understanding obstructive sleep apnea. Resp Physiol Neurobi. 2003;135(2):187–205.

23. Liang J, Cade BE, Wang H, Chen H, Gleason KJ, Larkin EK, et al. Comparison of Heritability Estimation and Linkage Analysis for Multiple Traits Using Principal Component Analyses. Genet Epidemiol. 2016;40(3):222–32.

24. Cade BE, Chen H, Stilp AM, Louie T, Ancoli-Israel S, Arens R, et al. Associations of variants In the hexokinase 1 and interleukin 18 receptor regions with oxyhemoglobin saturation during sleep. Plos Genet. 2019;15(4):e1007739.

25. Chen H, Cade BE, Gleason KJ, Bjonnes AC, Stilp AM, Sofer T, et al. Multiethnic Meta-Analysis Identifies \textitRAI1 as a Possible Obstructive Sleep Apnea–related Quantitative Trait Locus in Men. Am J Resp Cell Mol. 2018;58(3):391–401.

26. Wang H, Cade BE, Sofer T, Sands SA, Chen H, Browning SR, et al. Admixture mapping identifies novel loci for obstructive sleep apnea in Hispanic/Latino Americans. Hum Mol Genet. 2019;28(4):675–87.

27. Gusev A, Lee SH, Trynka G, Finucane H, Vilhjálmsson BJ, Xu H, et al. Partitioning Heritability of Regulatory and Cell-Type-Specific Variants across 11 Common Diseases. Am J Hum Genetics. 2014;95(5):535–52.

28. Maurano MT, Humbert R, Rynes E, Thurman RE, Haugen E, Wang H, et al. Systematic Localization of Common Disease-Associated Variation in Regulatory DNA. Science. 2012;337(6099):1190–5.

29. Trochet H, Pirinen M, Band G, Jostins L, McVean G, Spencer CCA. Bayesian meta-analysis across genome-wide association studies of diverse phenotypes. Genet Epidemiol. 2019;43(5):532–47.

30. Majumdar A, Haldar T, Bhattacharya S, Witte JS. An efficient Bayesian meta-analysis approach for studying cross-phenotype genetic associations. Plos Genet. 2018;14(2):e1007139.

31. Urbut SM, Wang G, Carbonetto P, Stephens M. Flexible statistical methods for estimating and testing effects in genomic studies with multiple conditions. Nat Genet. 2019;51(1):187–95.

32. Andreassen OA, Djurovic S, Thompson WK, Schork AJ, Kendler KS, O’Donovan MC, et al. Improved Detection of Common Variants Associated with Schizophrenia by Leveraging Pleiotropy with Cardiovascular-Disease Risk Factors. Am J Hum Genetics. 2013;92(2):197–209.

33. Andreassen OA, Thompson WK, Schork AJ, Ripke S, Mattingsdal M, Kelsoe JR, et al. Improved Detection of Common Variants Associated with Schizophrenia and Bipolar Disorder Using Pleiotropy-Informed Conditional False Discovery Rate. Plos Genet. 2013;9(4):e1003455.

34. Wang Y, Thompson WK, Schork AJ, Holland D, Chen CH, Bettella F, et al. Leveraging Genomic Annotations and Pleiotropic Enrichment for Improved Replication Rates in Schizophrenia GWAS. Plos Genet. 2016;12(1):e1005803.

35. Liley J, Wallace C. A Pleiotropy-Informed Bayesian False Discovery Rate Adapted to a Shared Control Design Finds New Disease Associations From GWAS Summary Statistics. Plos Genet. 2015;11(2):e1004926.

36. Giambartolomei C, Vukcevic D, Schadt EE, Franke L, Hingorani AD, Wallace C, et al. Bayesian Test for Colocalisation between Pairs of Genetic Association Studies Using Summary Statistics. Plos Genet. 2014;10(5):e1004383.

37. Giambartolomei C, Liu JZ, Zhang W, Hauberg M, Shi H, Boocock J, et al. A Bayesian framework for multiple trait colocalization from summary association statistics. Bioinformatics. 2018;34(15):2538–45.

38. Hormozdiari F, van de Bunt M, Segrè AV, Li X, Joo JWJ, Bilow M, et al. Colocalization of GWAS and eQTL Signals Detects Target Genes. Am J Hum Genetics. 2016;99(6):1245–60.

39. Chun S, Casparino A, Patsopoulos NA, Croteau-Chonka DC, Raby BA, Jager PLD, et al. Limited statistical evidence for shared genetic effects of eQTLs and autoimmune-disease-associated loci in three major immune-cell types. Nat Genet. 2017;49(4):600–5.

40. Zhu Z, Zhang F, Hu H, Bakshi A, Robinson MR, Powell JE, et al. Integration of summary data from GWAS and eQTL studies predicts complex trait gene targets. Nat Genet. 2016;48(5):481–7.

41. Gusev A, Ko A, Shi H, Bhatia G, Chung W, Penninx BWJH, et al. Integrative approaches for large-scale transcriptome-wide association studies. Nat Genet. 2016;48(3):245–52.

42. Larkin EK, Patel SR, Zhu X, Tracy RP, Jenny NS, Reiner AP, et al. A Study of The Relationship between The Interleukin-6 Gene and Obstructive Sleep Apnea. Clin Transl Sci. 2010;3(6):337–9.

43. Larkin EK, Patel SR, Goodloe RJ, Li Y, Zhu X, Gray-McGuire C, et al. A candidate gene study of obstructive sleep apnea in European Americans and African Americans. Am J Resp Crit Care. 2010;182(7):947–53.

44. Patel SR, Goodloe R, De G, Kowgier M, Weng J, Buxbaum SG, et al. Association of genetic loci with sleep apnea in European Americans and African-Americans: the Candidate Gene Association Resource (CARe). Plos One. 2012;7(11):e48836.

45. Geovanini GR, Wang R, Weng J, Tracy R, Jenny NS, Goldberger AL, et al. Elevations in neutrophils with obstructive sleep apnea: The Multi-Ethnic Study of Atherosclerosis (MESA). Int J Cardiol. 2018;257:318–23.

46. Geovanini GR, Wang R, Weng J, Jenny NS, Shea S, Allison M, et al. Association between Obstructive Sleep Apnea and Cardiovascular Risk Factors: Variation by Age, Sex, and Race. The Multi-Ethnic Study of Atherosclerosis. Ann Am Thorac Soc. 2018;15(8):970–7.

47. Kim JS, Podolanczuk AJ, Borker P, Kawut SM, Raghu G, Kaufman JD, et al. Obstructive Sleep Apnea and Subclinical Interstitial Lung Disease in the Multi-Ethnic Study of Atherosclerosis (MESA). Ann Am Thorac Soc. 2017;14(12):1786–95.

48. Lederer DJ, Jelic S, Basner RC, Ishizaka A, Bhattacharya J. Circulating KL-6, a Biomarker of Lung Injury, in Obstructive Sleep Apnea. Eur Respir J. 2009;33(4):793–6.

49. McNicholas WT. Comorbid obstructive sleep apnoea and chronic obstructive pulmonary disease and the risk of cardiovascular disease. Journal of Thoracic Disease. 2018;1(1):S4253–61–S4261.

50. McAlpine CS, Kiss MG, Rattik S, He S, Vassalli A, Valet C, et al. Sleep modulates haematopoiesis and protects against atherosclerosis. Nature. 2019;566(7744):383–7.

51. Xie J, Li F, Wu X, Hou W. Prevalence of pulmonary embolism in patients with obstructive sleep apnea and chronic obstructive pulmonary disease: The overlap syndrome. Heart Lung. 2019;48(3):261–5.

52. Conneely KN, Boehnke M. So Many Correlated Tests, So Little Time! Rapid Adjustment of P Values for Multiple Correlated Tests. Am J Hum Genetics. 2007;81(6):1158–68.

53. Boyle EA, Li YI, Pritchard JK. An Expanded View of Complex Traits: From Polygenic to Omnigenic. Cell. 2017;169(7):1177–86.

54. Butler MP, Emch JT, Rueschman M, Sands SA, Shea SA, Wellman A, et al. Apnea-Hypopnea Event Duration Predicts Mortality in Men and Women in the Sleep Heart Health Study. Am J Resp Crit Care. 2019;199(7):903–12.

55. Astle WJ, Elding H, Jiang T, Allen D, Ruklisa D, Mann AL, et al. The Allelic Landscape of Human Blood Cell Trait Variation and Links to Common Complex Disease. Cell. 2016;167(5):1415–1429.e19.

56. Loh PR, Kichaev G, Gazal S, Schoech AP, Price AL. Mixed-model association for biobank-scale datasets. Nat Genet. 2018;50(7):906–8.

57. Bycroft C, Freeman C, Petkova D, Band G, Elliott LT, Sharp K, et al. The UK Biobank resource with deep phenotyping and genomic data. Nature. 2018;562(7726):203–9.

58. Iotchkova V, Huang J, Morris JA, Jain D, Barbieri C, Walter K, et al. Discovery and refinement of genetic loci associated with cardiometabolic risk using dense imputation maps. Nat Genet. 2016;48(11):1303–12.

59. Hemani G, Zheng J, Elsworth B, Wade KH, Haberland V, Baird D, et al. The MR-Base platform supports systematic causal inference across the human phenome. Elife. 2018;7:e34408.

60. Gros A, Ollivier V, Ho-Tin-Noé B. Platelets in Inflammation: Regulation of Leukocyte Activities and Vascular Repair. Front Immunol. 2015;5:678.

61. Kichaev G, Bhatia G, Loh PR, Gazal S, Burch K, Freund MK, et al. Leveraging Polygenic Functional Enrichment to Improve GWAS Power. Am J Hum Genetics. 2019;104(1):65–75.

62. Finucane HK, Bulik-Sullivan B, Gusev A, Trynka G, Reshef Y, Loh PR, et al. Partitioning heritability by functional annotation using genome-wide association summary statistics. Nat Genet. 2015;47(11):1228–35.

63. Consortium TGte. The GTEx Consortium atlas of genetic regulatory effects across human tissues. Science. 2020;369(6509):1318–30.

64. Chen L, Ge B, Casale FP, Vasquez L, Kwan T, Garrido-Martín D, et al. Genetic Drivers of Epigenetic and Transcriptional Variation in Human Immune Cells. Cell. 2016;167(5):1398–1414.e24.

65. Owens RL, Edwards BA, Eckert DJ, Jordan AS, Sands SA, Malhotra A, et al. An Integrative Model of Physiological Traits Can be Used to Predict Obstructive Sleep Apnea and Response to Non Positive Airway Pressure Therapy. Sleep. 2015;38(6):961–70.

66. Huang T, Goodman M, Li X, Sands SA, Li J, Stampfer MJ, et al. C-reactive Protein and Risk of OSA in Four US Cohorts. Chest. 2021;159(6):2439–48.

67. Wijk R van, Solinge WW van. The energy-less red blood cell is lost: erythrocyte enzyme abnormalities of glycolysis. Blood. 2005;106(13):4034–42.

68. Powell FL, Fu Z. HIF-1 and ventilatory acclimatization to chronic hypoxia. Resp Physiol Neurobi. 2008;164(1–2):282–7.

69. Devinney MJ, Nichols NL, Mitchell GS. Sustained Hypoxia Elicits Competing Spinal Mechanisms of Phrenic Motor Facilitation. J Neurosci. 2016;36(30):7877–85.

70. Parry DA, Tamayo-Orrego L, Carroll P, Marsh JA, Greene P, Murina O, et al. PRIM1 deficiency causes a distinctive primordial dwarfism syndrome. Gene Dev. 2020;34(21–22):1520–33.

71. Franczak A, Bil-Lula I, Sawicki G, Fenton M, Ayas N, Skomro R. Matrix metalloproteinases as possible biomarkers of obstructive sleep apnea severity – a systematic review. Sleep Med Rev. 2019;46:9–16.

72. Dantas DA da S, Mauad T, Silva LFF, Lorenzi-Filho G, Formigoni GGS, Cahali MB. The Extracellular Matrix of the Lateral Pharyngeal Wall in Obstructive Sleep Apnea. Sleep. 2012;35(4):483–90.

73. Rosa DP da, Forgiarini LF, Baronio D, Feijó CA, Martinez D, Marroni NP. Simulating Sleep Apnea by Exposure to Intermittent Hypoxia Induces Inflammation in the Lung and Liver. Mediat Inflamm. 2012;2012:879419.

74. Zhu S, Zhou Y, Wang L, Zhang J, Wu H, Xiong J, et al. Transcriptional upregulation of MT2-MMP in response to hypoxia is promoted by HIF-1α in cancer cells. Mol Carcinogen. 2011;50(10):770–80.

75. García-Alvarez J, Ramirez R, Sampieri CL, Nuttall RK, Edwards DR, Selman M, et al. Membrane type-matrix metalloproteinases in idiopathic pulmonary fibrosis. Sarcoidosis Vasc Diffuse Lung Dis Official J Wasog World Assoc Sarcoidosis Other Granulomatous Disord. 2006;23(1):13–21.

76. Owens RL, Malhotra A. Sleep-disordered breathing and COPD: the overlap syndrome. Respir Care. 2010;55(10):1333–44; discussion 1344-6.

77. Yu LM, Zhang WH, Han XX, Li YY, Lu Y, Pan J, et al. Hypoxia-Induced ROS Contribute to Myoblast Pyroptosis during Obstructive Sleep Apnea via the NF-κB/HIF-1α Signaling Pathway. Oxid Med Cell Longev. 2019;2019:4596368.

78. Estephan E de P, Sobreira CF da R, Santos ACJD, Tomaselli PJ, Marques W, Ortega RPM, et al. A common CHRNE mutation in Brazilian patients with congenital myasthenic syndrome. J Neurol. 2018;265(3):708–13.

79. The Atherosclerosis Risk in Communities (ARIC) Study: design and objectives. The ARIC investigators. Am J Epidemiol. 1989;129(4):687–702.

80. Orwoll E, Blank JB, Barrett-Connor E, Cauley J, Cummings S, Ensrud K, et al. Design and baseline characteristics of the osteoporotic fractures in men (MrOS) study--a large observational study of the determinants of fracture in older men. Contemp Clin Trials. 2005;26(5):569–85.

81. Bild DE, Bluemke DA, Burke GL, Detrano R, Roux AVD, Folsom AR, et al. Multi-Ethnic Study of Atherosclerosis: objectives and design. Am J Epidemiol. 2002;156(9):871–81.

82. Fried LP, Borhani NO, Enright P, Furberg CD, Gardin JM, Kronmal RA, et al. The Cardiovascular Health Study: design and rationale. Ann Epidemiol. 1991;1(3):263–76.

83. Mukherjee S, Hillman D, Lee J, Fedson A, Simpson L, Ward K, et al. Cohort profile: the Western Australian Sleep Health Study. Sleep Breath. 2012;16(1):205–15.

84. Quan SF, Howard BV, Iber C, Kiley JP, Nieto FJ, O’Connor GT, et al. The Sleep Heart Health Study: design, rationale, and methods. Sleep. 1997;20(12):1077–85.

85. Chen X, Wang R, Zee P, Lutsey PL, Javaheri S, Alcántara C, et al. Racial/Ethnic Differences in Sleep Disturbances: The Multi-Ethnic Study of Atherosclerosis (MESA). Sleep. 2015;38(6):877– 88.

86. Blank JB, Cawthon PM, Carrion-Petersen ML, Harper L, Johnson JP, Mitson E, et al. Overview of recruitment for the osteoporotic fractures in men study (MrOS). Contemp Clin Trials. 2005;26(5):557–68.

87. Mehra R, Stone KL, Blackwell T, Israel SA, Dam TTL, Stefanick ML, et al. Prevalence and Correlates of Sleep-Disordered Breathing in Older Men: Osteoporotic Fractures in Men Sleep Study. J Am Geriatr Soc. 2007;55(9):1356–64.

88. Sorlie PD, Avilés-Santa LM, Wassertheil-Smoller S, Kaplan RC, Daviglus ML, Giachello AL, et al. Design and implementation of the Hispanic Community Health Study/Study of Latinos. Ann Epidemiol. 2010;20(8):629–41.

89. Redline S, Sotres-Alvarez D, Loredo J, Hall M, Patel SR, Ramos A, et al. Sleep-disordered Breathing in Hispanic/Latino Individuals of Diverse Backgrounds. The Hispanic Community Health Study/Study of Latinos. Am J Resp Crit Care. 2014;189(3):335–44.

90. Hanis CL, Ferrell RE, Barton SA, Aguilar L, Garza-Ibarra A, Tulloch BR, et al. Diabetes among Mexican Americans in Starr County, Texas. Am J Epidemiol. 1983;118(5):659–72.

91. Redline S, Tishler PV, Tosteson TD, Williamson J, Kump K, Browner I, et al. The familial aggregation of obstructive sleep apnea. Am J Resp Crit Care. 1995;151(3 Pt 1):682–7.

92. Feinleib M. The Framingham Study: sample selection, follow-up, and methods of analyses. National Cancer Inst Monogr. 1985;67:59–64.

93. Hanis CL, Redline S, Cade BE, Bell GI, Cox NJ, Below JE, et al. Beyond type 2 diabetes, obesity and hypertension: an axis including sleep apnea, left ventricular hypertrophy, endothelial dysfunction, and aortic stiffness among Mexican Americans in Starr County, Texas. Cardiovasc Diabetol. 2016;15(1):86.

94. Sudlow C, Gallacher J, Allen N, Beral V, Burton P, Danesh J, et al. UK Biobank: An Open Access Resource for Identifying the Causes of a Wide Range of Complex Diseases of Middle and Old Age. Plos Med. 2015;12(3):e1001779.

95. Kerimov N, Hayhurst JD, Peikova K, Manning JR, Walter P, Kolberg L, et al. A compendium of uniformly processed human gene expression and splicing quantitative trait loci. Nat Genet. 2021;53(9):1290–9.

