## Supplementary material for "Leveraging pleiotropy to discover and interpret GWAS results for sleep-associated traits": Revision Supplementary Figures

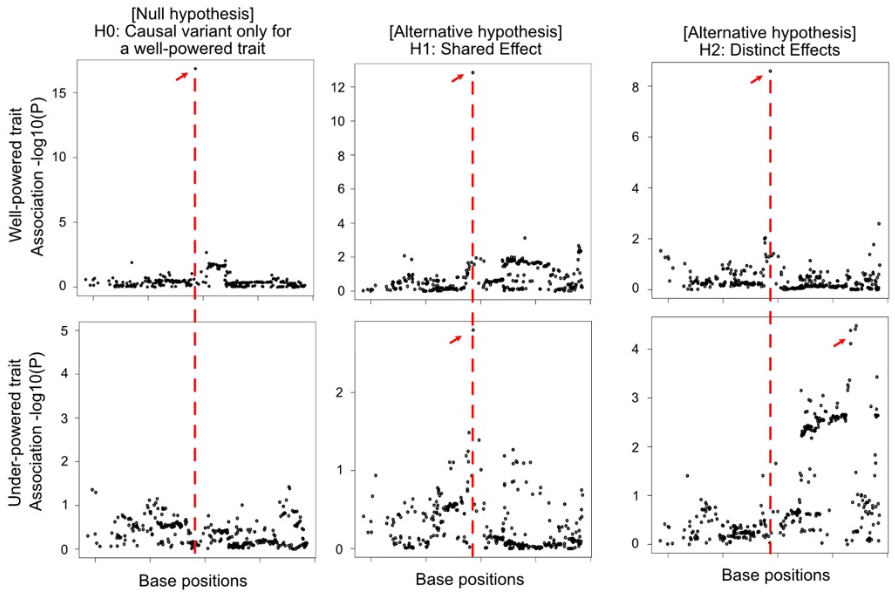


**Figure S1. Schematic of JLIM analysis.** Examples of trait pairs simulated under the null - no causal effect for an underpowered trait ($H_0$), shared effect between well-powered and underpowered traits ($H_1$), and distinct effects between two compared traits ($H_2$). $H_1$ and $H_2$ are competing alternative hypotheses. The simulated true causal variants are indicated by red arrows.

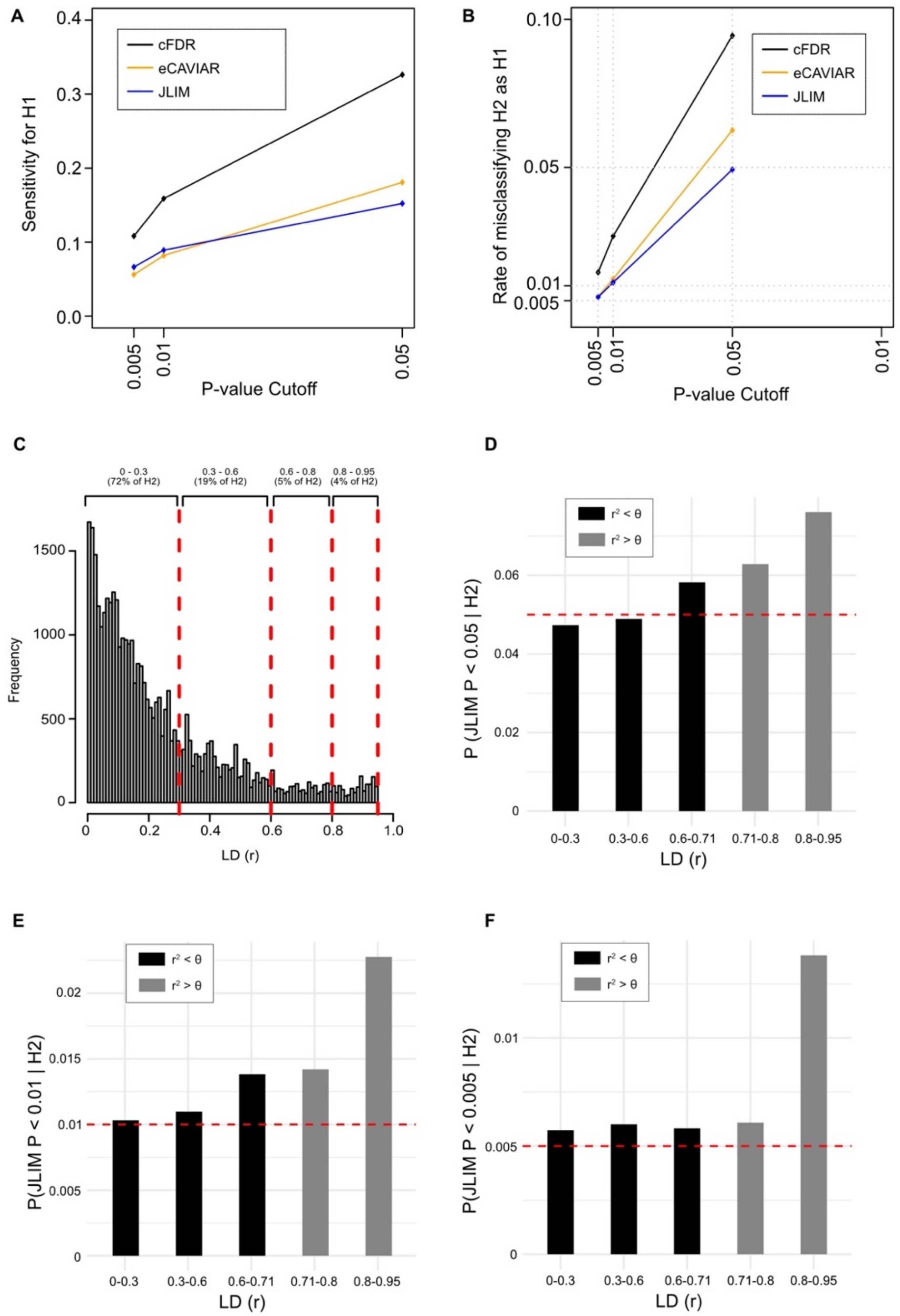


**Figure S2. Sensitivity for $H_1$ and specificity to distinguish $H_1$ from $H_2$. (A)** Sensitivity to detect $H_1$ at the same false positive rate for $H_0$. The same p-value cutoff was used for JLIM and cFDR. For eCAVIAR, a posterior cutoff was calibrated to match the P-value cutoff using unfiltered $H_0$ simulation data. **(B)** The rate of misclassifying $H_2$ as $H_1$. $H_2$ loci include all loci simulating distinct causative variants in LD between 0 to 0.95. Again, the cutoffs of cFDR, eCAVIAR and JLIM were calibrated to the same specificity using $H_0$. **(C)** The distribution of LD for randomly selected pairs of SNPs. This distribution has been drawn from the LD patterns between random pairs of SNPs within 80 random loci (200kb each) in the population of European ancestry. This distribution was used to simulate the LD between distinct causative variants in $H_2$. **(D,E,F)** The rate of misclassifying $H_2$ as $H_1$, broken down by the LD between simulated distinct causative variants for two traits. The cutoff of JLIM P-values was set to **(D)** 0.05, **(E)** 0.01 or **(F)** 0.005. The dashed horizontal line indicates the JLIM P-value cutoff. θ represents the genetic resolution parameter for JLIM, set to 0.5 in this study. JLIM does not claim to distinguish $H_2$ beyond the specified genetic resolution limit ($r^2 > \theta$; light grey bars).

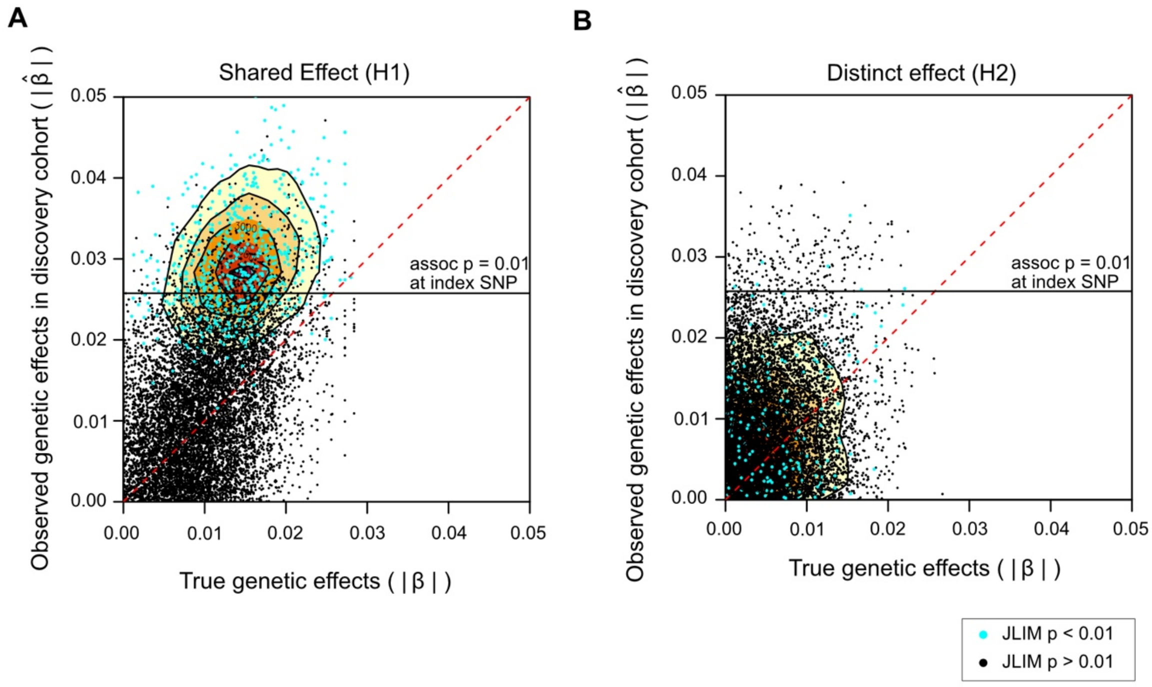


**Figure S3. Winner's curse.** The true and observed genetic effect sizes for underpowered traits are shown to highlight the winner's curse in a discovery cohort. We show data from 10,000 loci each simulated under **(A)** $H_1$ and **(B)** $H_2$. In all panels, each dot represents the genetic effects measured at the index SNPs. The index SNPs are defined as the lead SNPs of association to well-powered clinical traits. Occasionally, the index SNPs deviate from the causative SNPs due to sampling noise, and when this happens, we calculated the true effect size at the index SNP by multiplying the true effect of the causative SNP by the LD between the index and causative SNPs. The loci detected at JLIM P < 0.01 are indicated by cyan dots, and their density distribution are shown in contours. The loci found by cFDR (association $P < 5x10^{-8}$ for well-powered trait and < 0.01 for underpowered trait) are represented by dots above the black horizontal line.

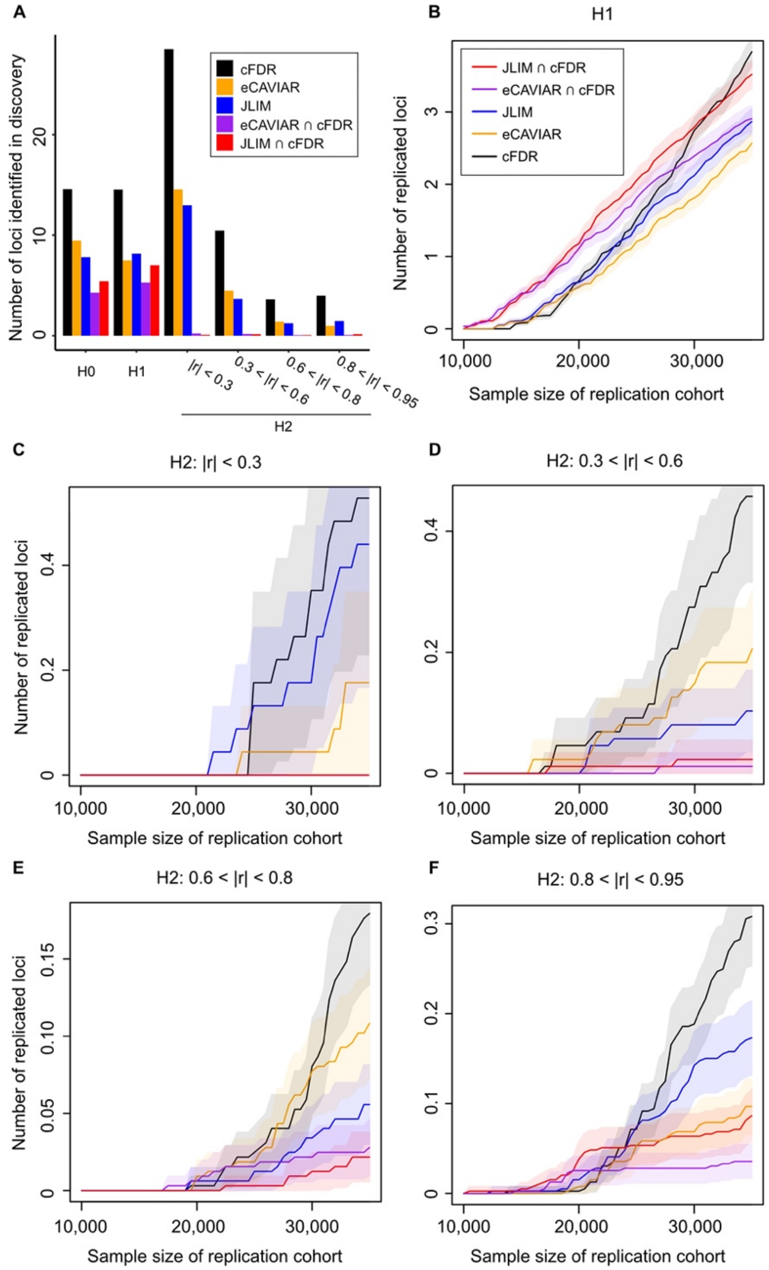


**Figure S4. The number of loci that were identified in simulated discovery and validation cohorts, broken down by the configuration of causative variants. (A)** The number of loci identified in a discovery cohort by pleiotropy analysis. **(B-F)** The number of loci replicated in a validation cohort, subdivided by the configuration of causative variants: **(B)** $H_1$, $H_2$ with the LD between causative variants to be in the ranges of **(C)** $|r| < 0.3$, **(D)** $0.3 < |r| < 0.6$ , **(E)** $0.6 < |r| < 0.8$, and **(F)** $0.8 < |r| < 0.95$. In all panels, simulation was conducted under the following parameters: A total of 2,500 association peaks from well-powered GWAS studies (n=150,000) were tested for pleiotropy in simulated discovery cohorts (n=10,000), and then the candidate pleiotropic loci were tested for replication in simulated validation cohorts of the same genetic ancestry (n=10,000-35,000). The candidate loci were identified by conditional false discovery rate (cFDR), eCAVIAR, Joint Likelihood Mapping (JLIM), or the intersection of eCAVIAR or JLIM and cFDR, all at the P-value cutoff of 0.01 (or equivalent posterior cutoff). The 2,500 GWAS peaks consist of the loci simulating no causal effect for underpowered traits ($H_0$) and those simulating the same causal effect between two traits ($H_1$) or distinct causal effects ($H_2$). The proportion of $H_0$ was set to 30%, and the remaining 70% of loci were split to $H_1$ and $H_2$ at the ratio of 1:19. The effect sizes of causative variants are correlated ($\rho = 0.7$) under $H_1$ but uncorrelated under $H_2$. Bonferroni correction was applied on replication tests. The shaded area denotes the 95% CIs.

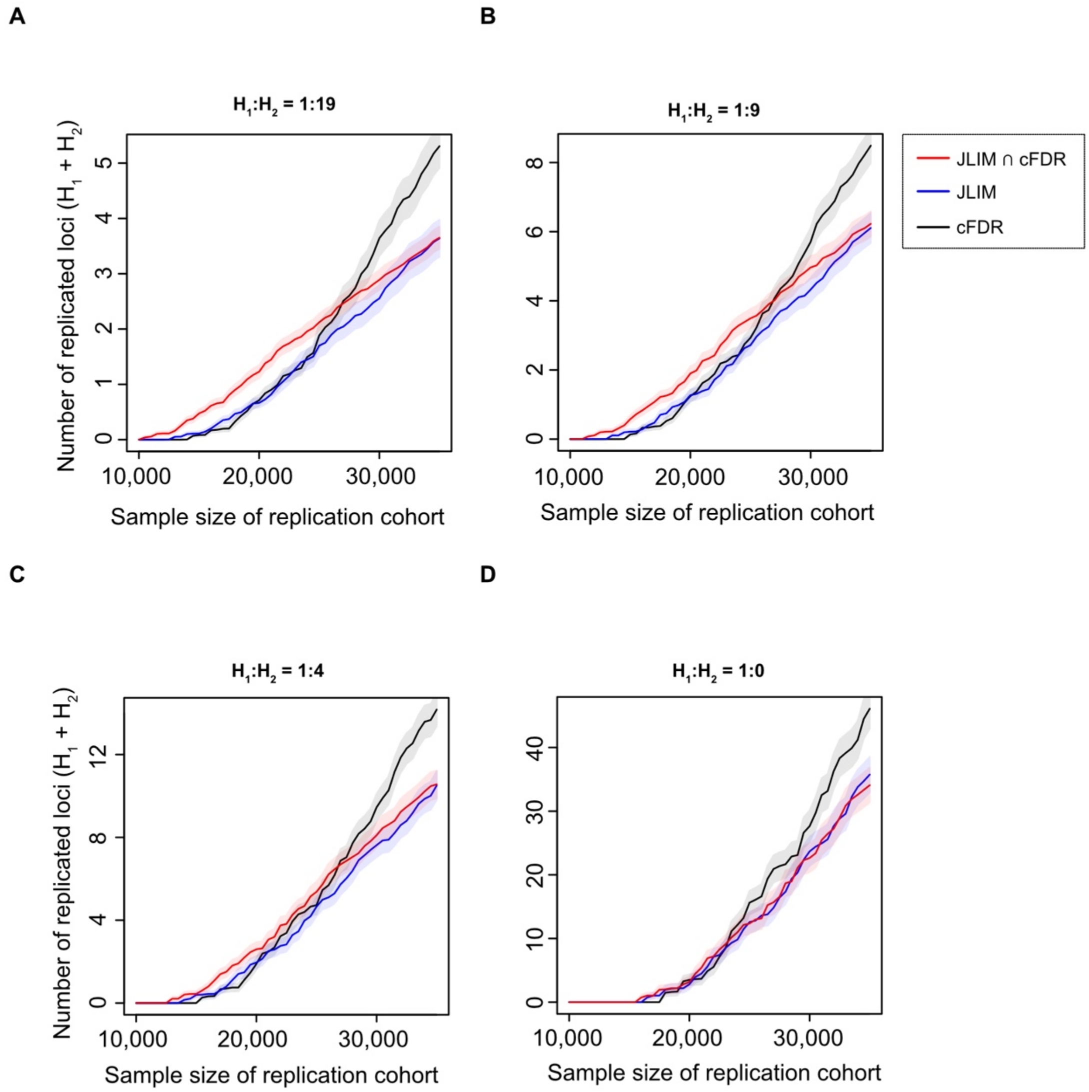


**Figure S5. The projected number of replicated loci in simulations, varying the relative ratio between $H_1$ and $H_2$.** A total of 2,500 association peaks from well-powered GWAS studies (n=150,000) were tested for pleiotropy in a discovery cohort (n=10,000), and then the candidate pleiotropic loci were tested for replication in an independent validation cohort of the same genetic ancestry (n=10,000-35,000). The candidate loci were identified by conditional false discovery rate (cFDR), Joint Likelihood Mapping (JLIM), or the intersection of both, all at the P-value cutoff of 0.01. The 2,500 GWAS peaks consist of the loci simulating no causal effect for underpowered traits ($H_0$) and those simulating the same causal effect between two traits ($H_1$) or distinct causal effects ($H_2$). The proportion of $H_0$ was set to 30%, and the remaining 70% of loci were split to $H_1$ and $H_2$ at the ratio of **(A)** 1:19, **(B)** 1:9, **(C)** 1:4, and **(D)** 1:0. The effect sizes of causative variants are correlated ($\rho = 0.7$) under $H_1$ but uncorrelated under $H_2$. Bonferroni correction was applied on replication tests. The shaded area denotes the 95% CIs.

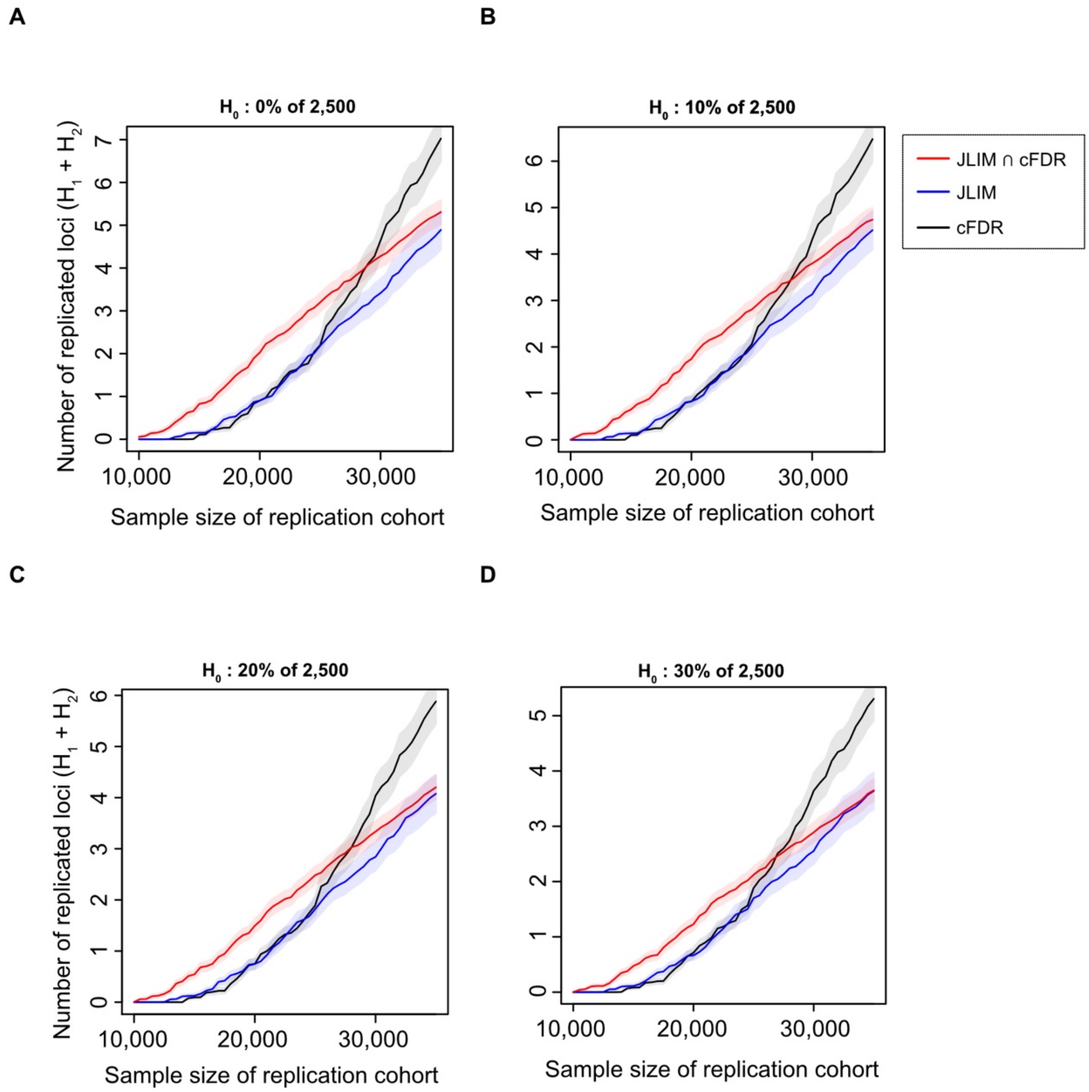


**Figure S6. The projected number of replicated loci in simulations, varying the proportion of $H_0$ loci.** A total of 2,500 association peaks from well-powered GWAS studies (n=150,000) were tested for pleiotropy in a discovery cohort (n=10,000), and then the candidate pleiotropic loci were tested for replication in an independent validation cohort of the same genetic ancestry (n=10,000-35,000). The candidate loci were identified by conditional false discovery rate (cFDR), Joint Likelihood Mapping (JLIM), or the intersection of both, all at the P-value cutoff of 0.01. The 2,500 GWAS peaks consist of the loci simulating no causal effect for underpowered traits ($H_0$) and those simulating the same causal effect between two traits ($H_1$) or distinct causal effects ($H_2$). The proportion of $H_0$ was varied to **(A)** 0%, **(B)** 10%, **(C)** 20%, and **(D)** 30%, and the remaining loci were split to $H_1$ and $H_2$ at the ratio of 1:19. The effect sizes of causative variants are correlated ($\rho$=0.7) under $H_1$ but uncorrelated under $H_2$. Bonferroni correction was applied on replication tests. The shaded area denotes the 95% CIs.

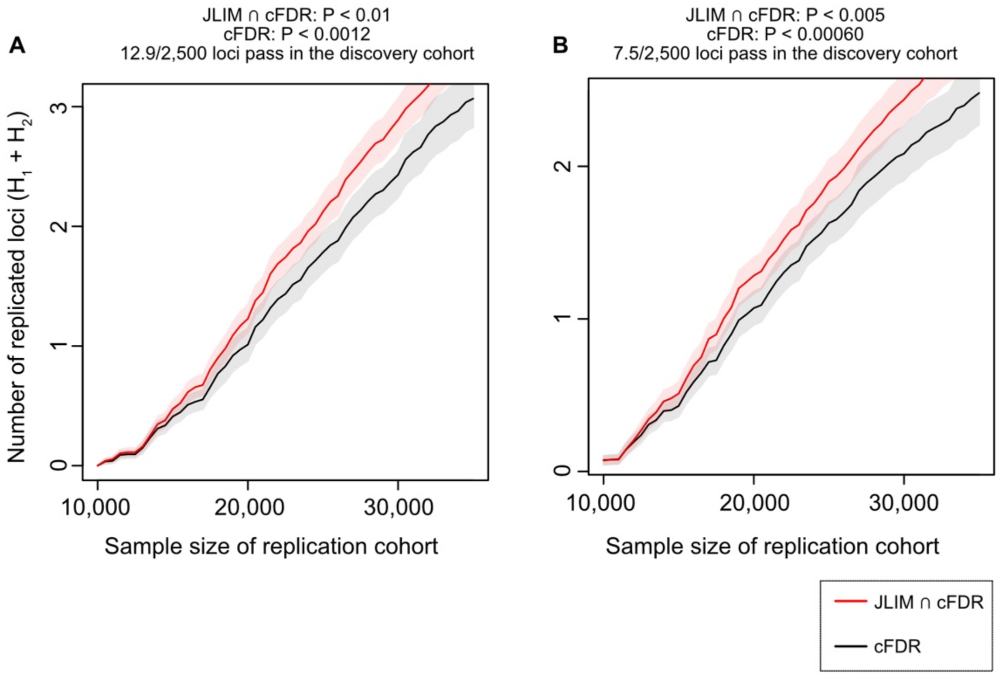


**Figure S7. The projected number of replicated loci in simulations where the cFDR threshold was tightened to identify the same number of candidate loci as the JLIM/cFDR consensus method in a discovery cohort.** A total of 2,500 association peaks from well-powered GWAS studies (n=150,000) were tested for pleiotropy in a discovery cohort (n=10,000), and then the candidate pleiotropic loci were tested for replication in an independent validation cohort of the same genetic ancestry (n=10,000-35,000). The candidate loci were identified by conditional false discovery rate (cFDR) or by the JLIM/cFDR consensus method. **(A)** The consensus method (red line) selected 12.9 candidate pleiotropic loci by taking the intersection between JLIM P < 0.01 and cFDR (association P < 0.01 for an underpowered trait). For the comparison, we tightened cFDR threshold to underpowered trait assoc P < 0.0012 so to identify the same number of candidate loci in a discovery cohort (black line). **(B)** Similarly, the consensus method (red line) found 7.5 candidate loci by taking the intersection between JLIM P < 0.005 and cFDR (assoc P < 0.005 for underpowered trait). The cFDR threshold was tightened to underpowered trait assoc P < 0.00060 for the same number of candidates (black line). In both panels, the 2,500 GWAS peaks consist of the loci simulating no causal effect for underpowered traits ($H_0$) and those simulating the same causal effect between two traits ($H_1$) or distinct causal effects ($H_2$). The proportion of $H_0$ was set to 30%, and the remaining 70% of loci were split to $H_1$ and $H_2$ at the ratio of 1:19. The effect sizes of causative variants are correlated (ρ=0.7) under $H_1$ but uncorrelated under $H_2$. Bonferroni correction was applied on replication tests. The shaded area denotes the 95% CIs.

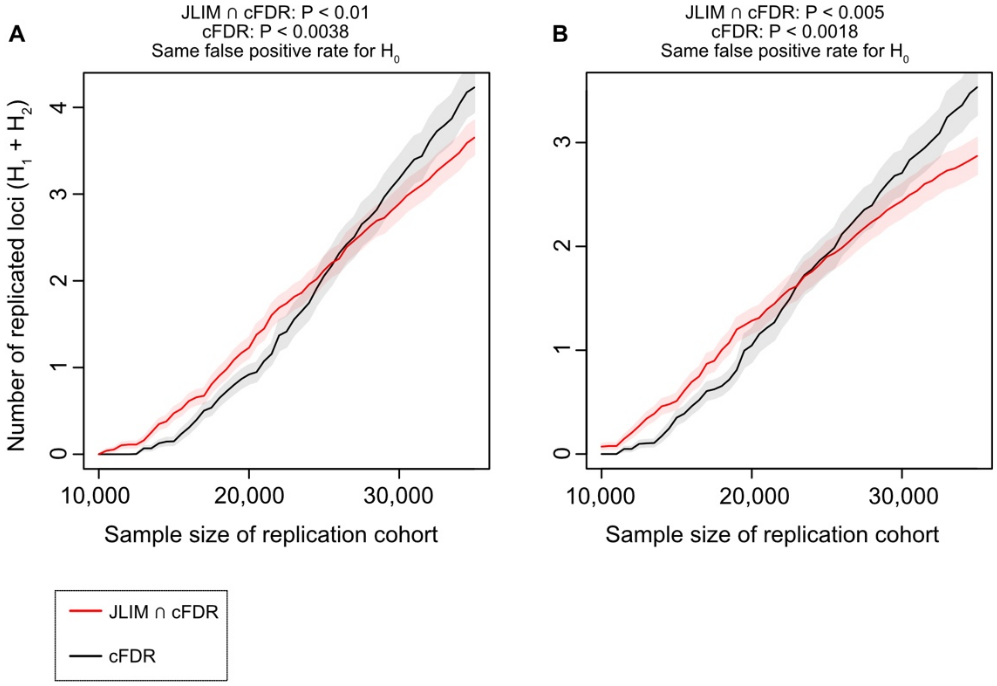


**Figure S8. The projected number of replicated loci in simulations where the cFDR threshold was calibrated to have the same false positive rate for $H_0$.** A total of 2,500 association peaks from well-powered GWAS studies (n=150,000) were tested for pleiotropy in a discovery cohort (n=10,000), and then the candidate pleiotropic loci were tested for replication in an independent validation cohort of the same genetic ancestry (n=10,000-35,000). The candidate loci were identified by conditional false discovery rate (cFDR) or by the JLIM/cFDR consensus method. **(A)** The consensus method (red line), by taking the intersection between JLIM P < 0.01 and cFDR P < 0.01, showed the empirical false positive rate of 0.0038 in simulated $H_0$ dataset. To match this false positive rate, we tightened cFDR threshold to P < 0.0038 (black line). **(B)** Similarly, the consensus method (red line) showed the empirical false positive rate of 0.0018 in $H_0$ when the intersection was taken at JLIM and cFDR P < 0.005. To match the false positive rate, the cFDR threshold was tightened to P < 0.0018 (black line). The cFDR P-value refers to the P-value of association to an underpowered trait. In both panels, the 2,500 GWAS peaks consist of the loci simulating no causal effect for underpowered traits ($H_0$) and those simulating the same causal effect between two traits ($H_1$) or distinct causal effects ($H_2$). The proportion of $H_0$ was set to 30%, and the remaining 70% of loci were split to $H_1$ and $H_2$ at the ratio of 1:19. The effect sizes of causative variants are correlated ($\rho$=0.7) under $H_1$ but uncorrelated under $H_2$. Bonferroni correction was applied on replication tests. The shaded area denotes the 95% CIs.

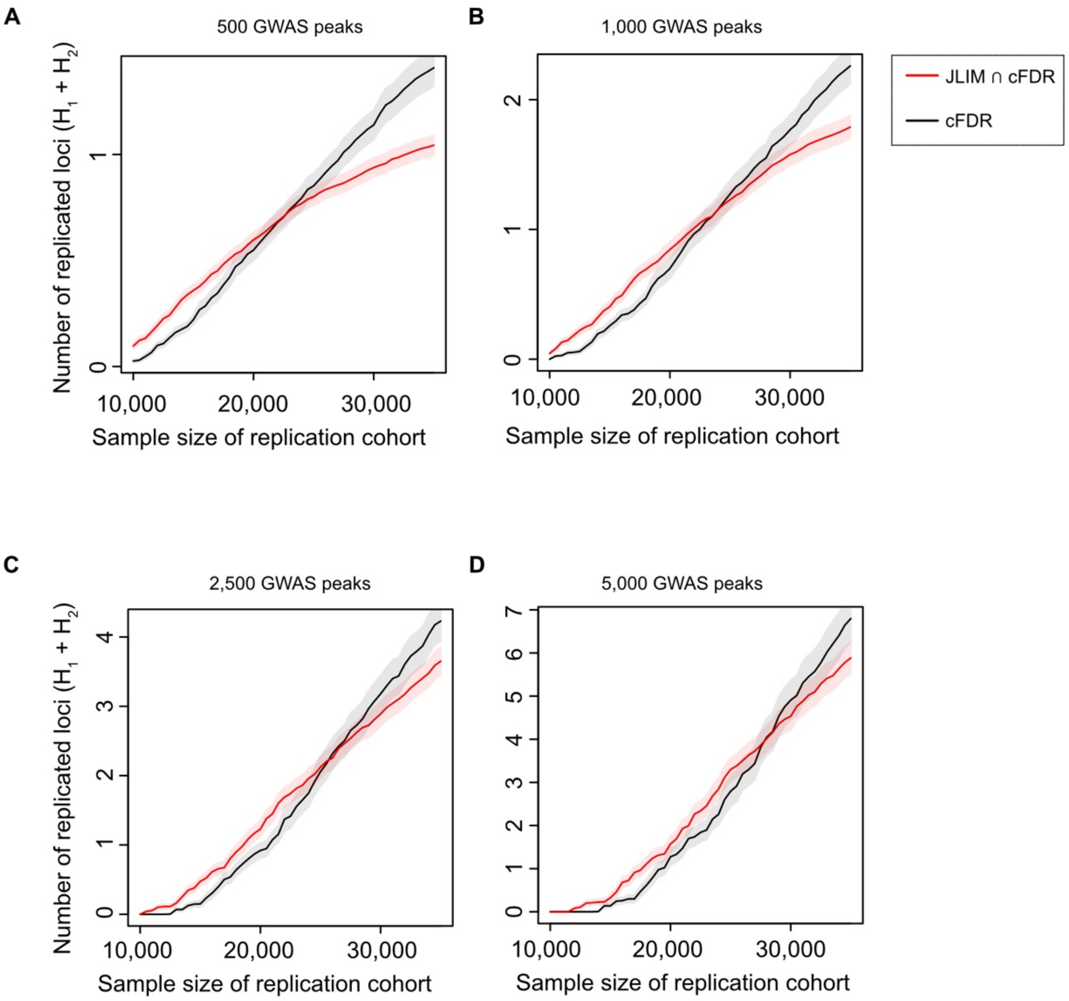


**Figure S9. The projected number of replicated loci in simulations where the number of association peaks available from well-powered traits varied.** A total of **(A)** 500, **(B)** 1,000, **(C)** 2,500, and **(D)** 5,000 association peaks from well-powered GWAS studies (n=150,000) were tested for pleiotropy in a discovery cohort (n=10,000), and then the candidate pleiotropic loci were tested for replication in an independent validation cohort of the same genetic ancestry (n=10,000-35,000). The candidate loci were identified by conditional false discovery rate (cFDR) or by the JLIM/cFDR consensus method. The consensus method (red line), by taking the intersection between JLIM P < 0.01 and cFDR P < 0.01, showed the empirical false positive rate of 0.0038 in simulated $H_0$ dataset. To match this false positive rate, we tightened cFDR threshold to P < 0.0038 (black line). The cFDR P-value refers to the P-value of association to an underpowered trait. The GWAS peaks consist of the loci simulating no causal effect for underpowered traits ($H_0$) and those simulating the same causal effect between two traits ($H_1$) or distinct causal effects ($H_2$). The proportion of $H_0$ was set to 30%, and the remaining 70% of loci were split to $H_1$ and $H_2$ at the ratio of 1:19. The effect sizes of causative variants are correlated ($\rho$=0.7) under $H_1$ but uncorrelated under $H_2$. Bonferroni correction was applied on replication tests. The shaded area denotes the 95% CIs.

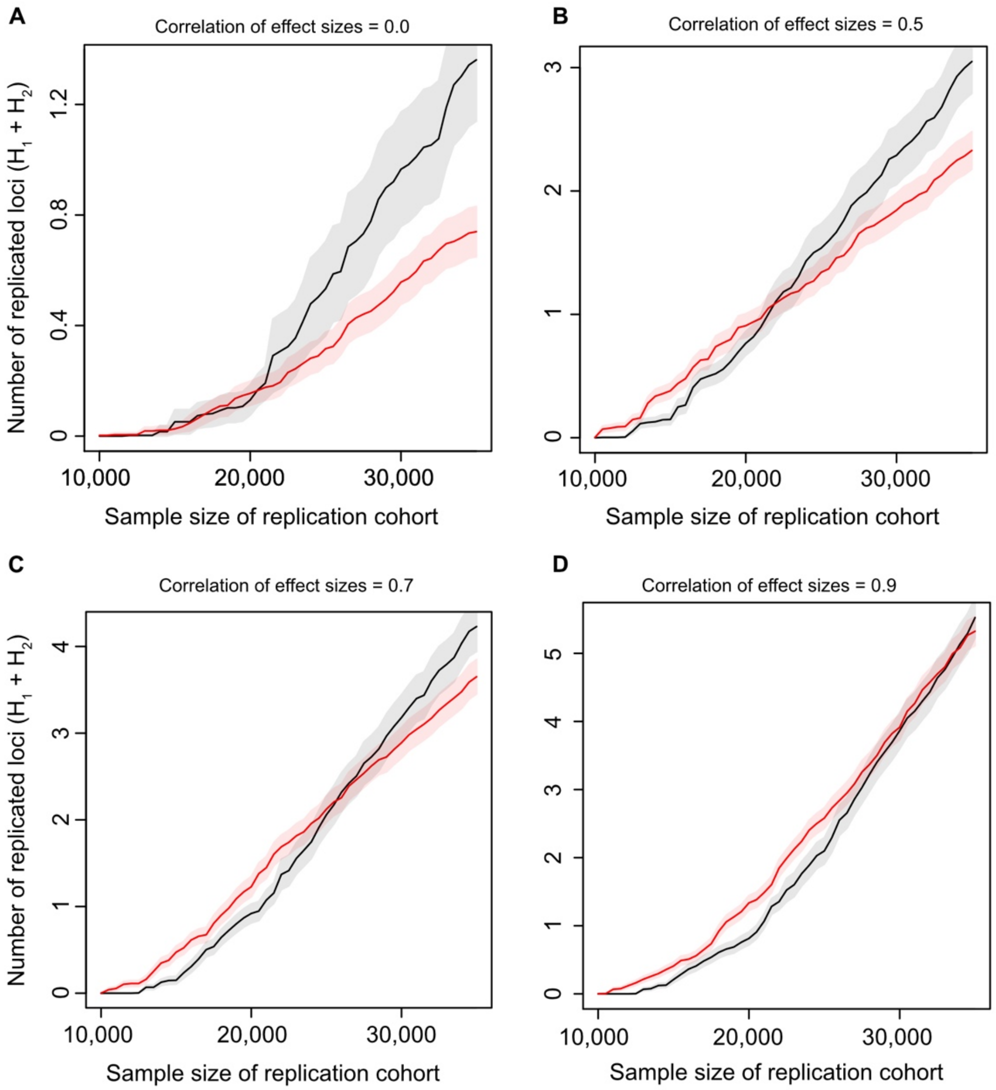


**Figure S10. The projected number of replicated loci in simulations where the correlation of effect sizes for $H_1$ varied.** A total of 2,500 association peaks from well-powered GWAS studies (n=150,000) were tested for pleiotropy in a discovery cohort (n=10,000), and then the candidate pleiotropic loci were tested for replication in an independent validation cohort of the same genetic ancestry (n=10,000-35,000). The candidate loci were identified by conditional false discovery rate (cFDR) or by the JLIM/cFDR consensus method. The consensus method (red line), by taking the intersection between JLIM $P < 0.01$ and cFDR $P < 0.01$, showed the empirical false positive rate of 0.0038 in simulated $H_0$ dataset. To match this false positive rate, we tightened cFDR threshold to $P < 0.0038$ (black line). The cFDR P-value refers to the P-value of association to an underpowered trait. The 2,500 GWAS peaks consist of the loci simulating no causal effect for underpowered traits ($H_0$) and those simulating the same causal effect between two traits ($H_1$) or distinct causal effects ($H_2$). The proportion of $H_0$ was set to 30%, and the remaining 70% of loci were split to $H_1$ and $H_2$ at the ratio of 1:19. The effect sizes of causative variants are correlated with **(A)** $\rho$ = 0.0, **(B)** $\rho$ = 0.5, **(C)** $\rho$ = 0.7 and **(D)** $\rho$ = 0.9 under $H_1$ but uncorrelated under $H_2$. In all panels, Bonferroni correction was applied on replication tests. The shaded area denotes the 95% CIs.

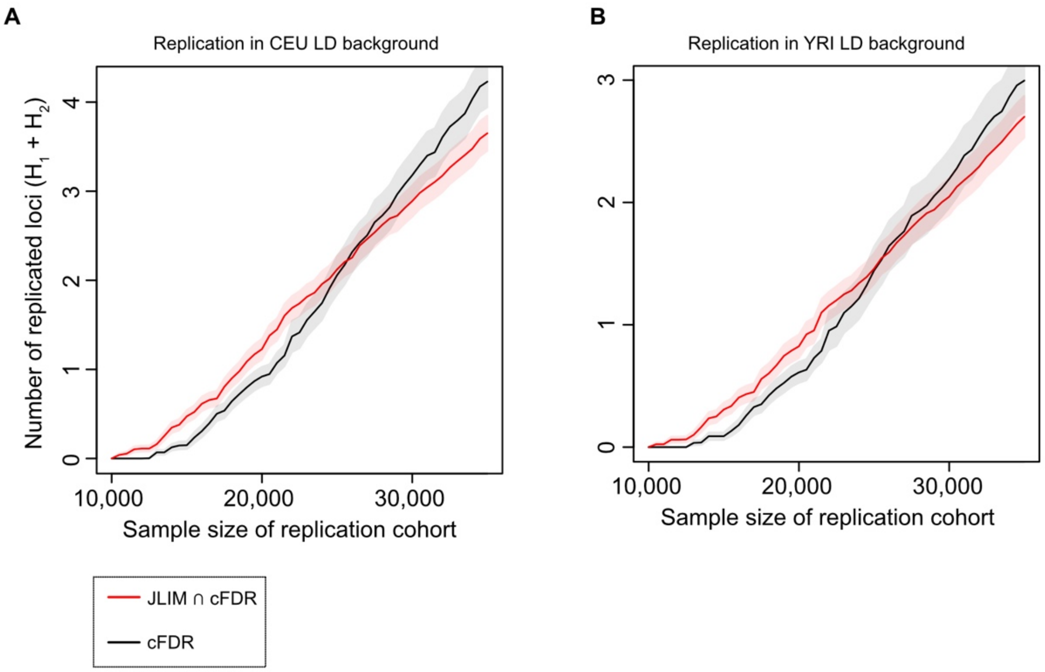


**Figure S11. The projected number of replicated loci in simulations of trans-ethnic replication.** A total of 2,500 association peaks from well-powered GWAS studies (n=150,000) were tested for pleiotropy in a discovery cohort (n=10,000), and then the candidate pleiotropic loci were tested for replication in an independent validation cohort of **(A)** the same (CEU) and **(B)** different (YRI) genetic ancestry (n=10,000-35,000). The LD patterns of genetic ancestry were obtained from the 1000 Genomes Project data. In both panels, the candidate loci were identified by conditional false discovery rate (cFDR) or by the JLIM/cFDR consensus method. The consensus method (red line), by taking the intersection between JLIM P < 0.01 and cFDR P < 0.01, showed the empirical false positive rate of 0.0038 in simulated $H_0$ dataset. To match this false positive rate, we tightened cFDR threshold to P < 0.0038 (black line). The cFDR P-value refers to the P-value of association to an underpowered trait. The 2,500 GWAS peaks consist of the loci simulating no causal effect for underpowered traits ($H_0$) and those simulating the same causal effect between two traits ($H_1$) or distinct causal effects ($H_2$). The proportion of $H_0$ was set to 30%, and the remaining 70% of loci were split to $H_1$ and $H_2$ at the ratio of 1:19. The effect sizes of causative variants are correlated with $\rho$=0.7 under $H_1$ but uncorrelated under $H_2$. Bonferroni correction was applied on replication tests. The shaded area denotes the 95% CIs.

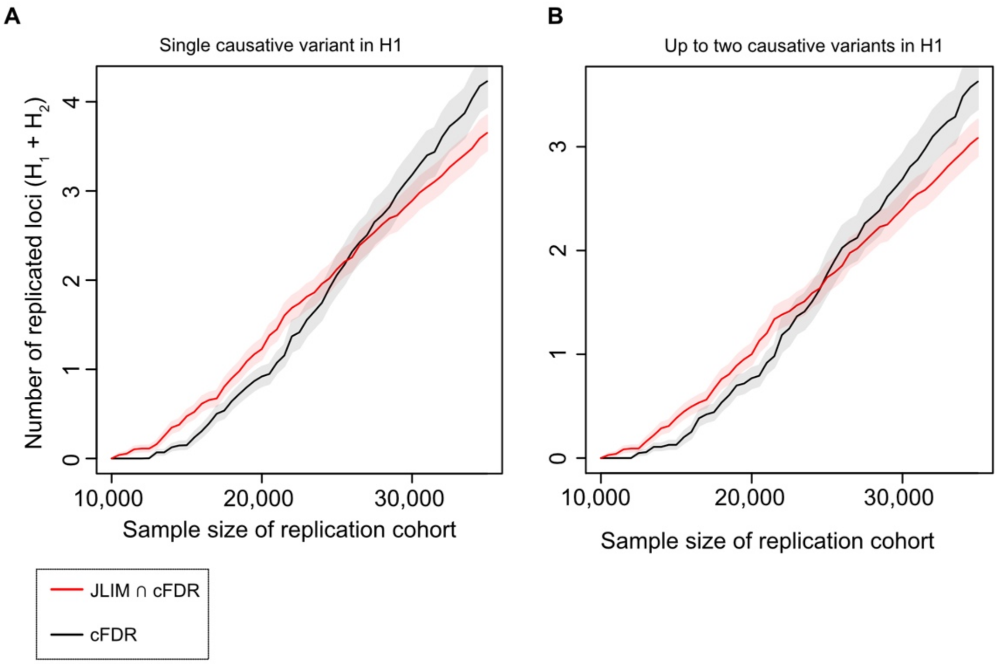


**Figure S12. The projected number of replicated loci in simulations of multiple causative variants for $H_1$.** A total of 2,500 association peaks from well-powered GWAS studies (n=150,000) were tested for pleiotropy in a discovery cohort (n=10,000), and then the candidate pleiotropic loci were tested for replication in an independent validation cohort of the same genetic ancestry (n=10,000-35,000). The candidate loci were identified by conditional false discovery rate (cFDR) or by the JLIM/cFDR consensus method. The consensus method (red line), by taking the intersection between JLIM $P < 0.01$ and cFDR $P < 0.01$, showed the empirical false positive rate of 0.0038 in simulated $H_0$ dataset. To match this false positive rate, we tightened cFDR threshold to $P < 0.0038$ (black line). The cFDR P-value refers to the P-value of association to an underpowered trait. The 2,500 GWAS peaks consist of the loci simulating no causal effect for underpowered traits ($H_0$) and those simulating the same causal effect between two traits ($H_1$) or distinct causal effects ($H_2$). The proportion of $H_0$ was set to 30%, and the remaining 70% of loci were split to $H_1$ and $H_2$ at the ratio of 1:19. In Panel **(A)**, only one causative variant was simulated for $H_1$, whereas in Panel **(B)**, up to two causative variants were simulated for $H_1$. The proportion of loci with two causative variants was set to 1/4 of all $H_1$ as expected under Poisson distribution with the causal fraction of 0.01. The effect sizes of causative variants are correlated with $\rho$=0.7 under $H_1$ but uncorrelated under $H_2$. Bonferroni correction was applied on replication tests. The shaded area denotes the 95% CIs.

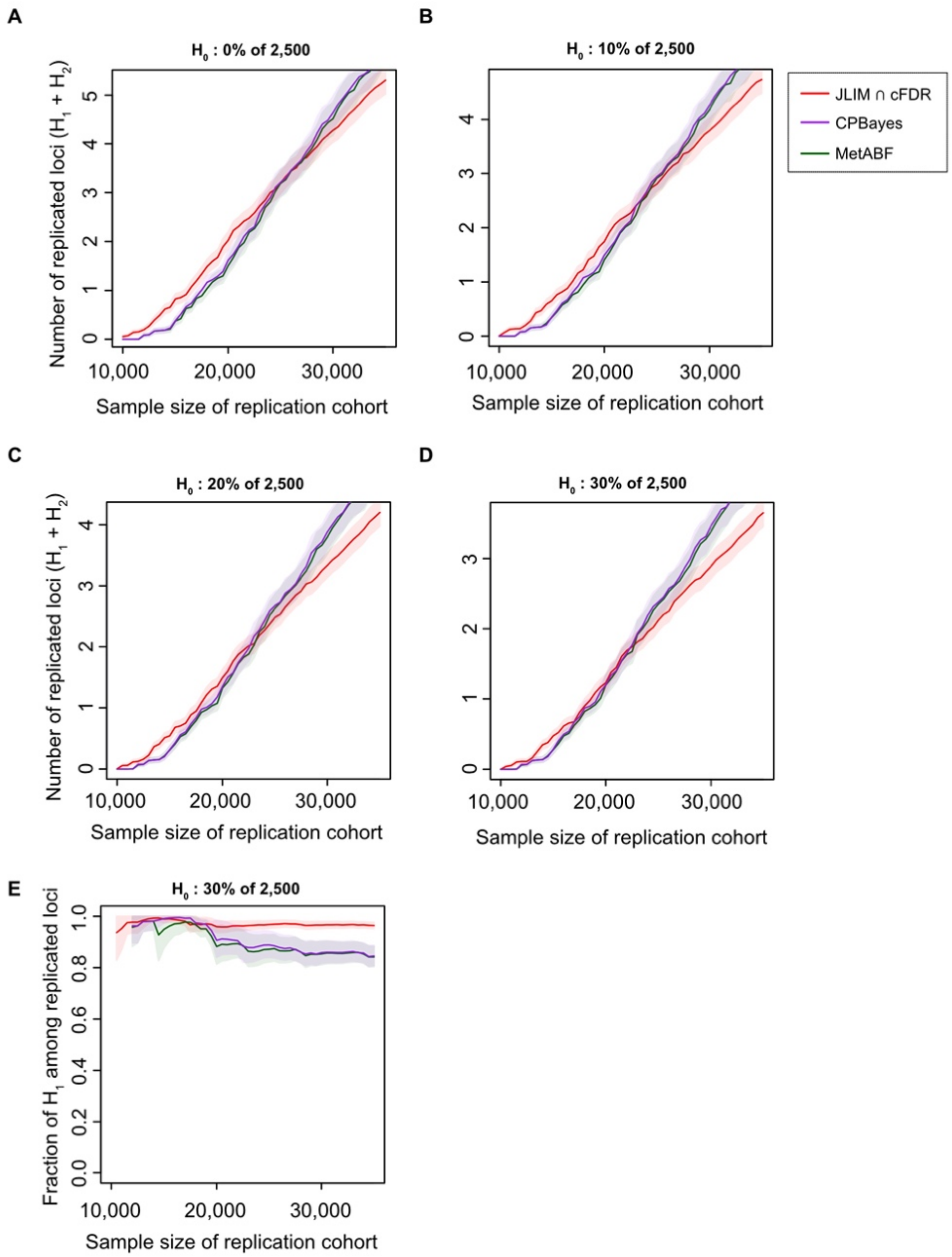


**Figure S13. Comparison with Bayesian meta-analysis methods with a validation cohort from the same ancestry.** A total of 2,500 association peaks from well-powered GWAS studies (n=150,000) were tested for pleiotropy in a discovery cohort (n=10,000), and then the candidate pleiotropic loci were tested for replication in an independent validation cohort of the same genetic ancestry (n=10,000-35,000). The candidate loci were identified by the JLIM/cFDR consensus method, MetABF and CPBayes. The consensus method (red line), by taking the intersection between JLIM $P < 0.01$ and cFDR $P < 0.01$, showed the empirical false positive rate of 0.0038 in simulated $H_0$ dataset. Bayesian posterior thresholds for MetABF (green) and CPBayes (purple) calibrated using $H_0$ loci to match the false positive rate of the consensus method. The 2,500 GWAS peaks consist of the loci simulating no causal effect for underpowered traits ($H_0$) and those simulating the same causal effect between two traits ($H_1$) or distinct causal effects ($H_2$). The proportion of $H_0$ varied to **(A)** 0%, **(B)** 10%, **(C)** 20% and **(D)** 30%, and the remaining loci were split to $H_1$ and $H_2$ at the ratio of 1:19. In all panels, the effect sizes of causative variants are correlated with $\rho=0.7$ under $H_1$ but uncorrelated under $H_2$. Bonferroni correction was applied on replication tests. The shaded area denotes the 95% CIs. In **(E)**, the fraction of $H_1$ among replicated loci is compared among three methods when the proportion of $H_0$ is 30% (similar for different proportions of $H_0$).

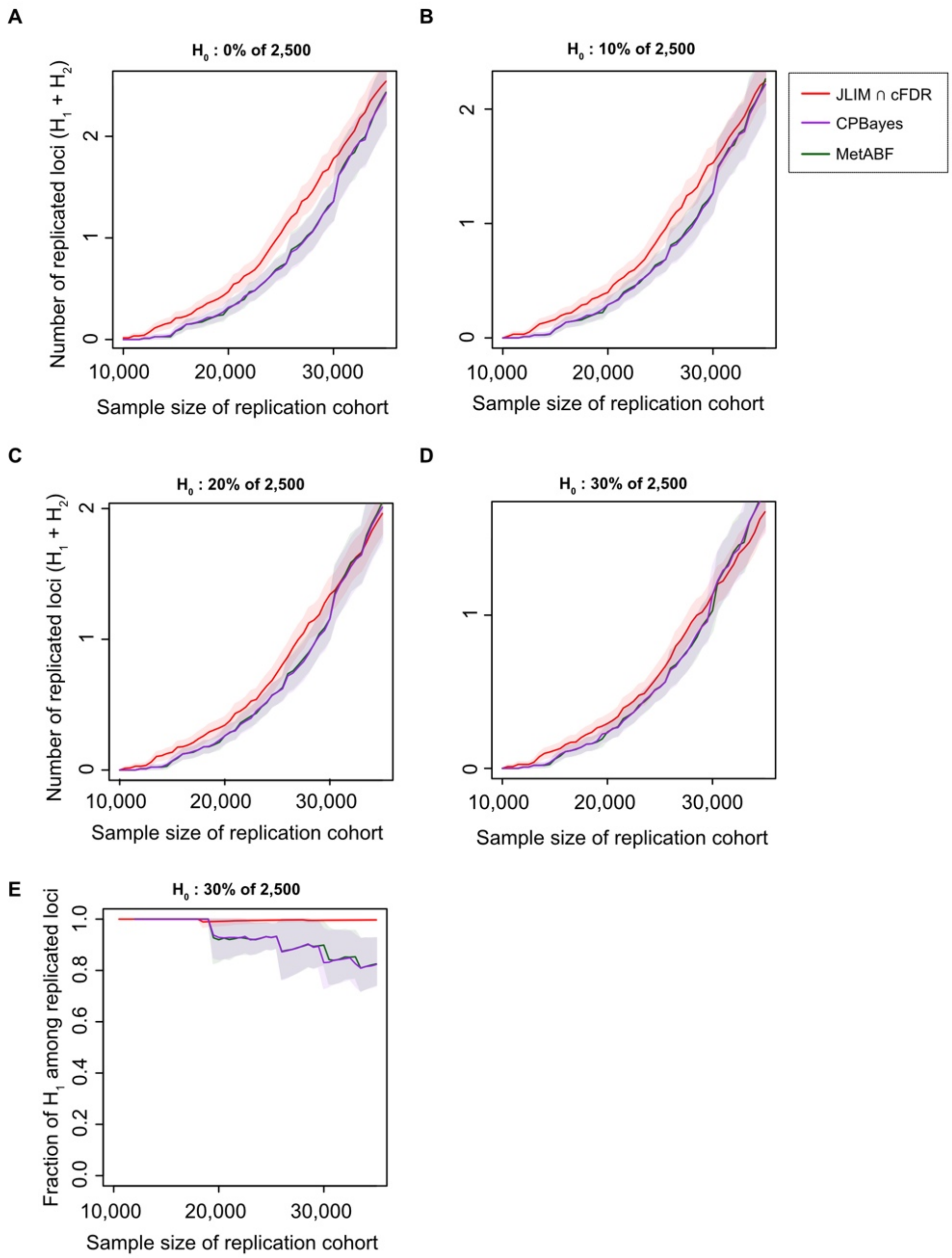


**Figure S14. Comparison with Bayesian meta-analysis methods with a validation cohort from the different ancestry (YRI).** A total of 2,500 association peaks from well-powered GWAS studies (n=150,000) were tested for pleiotropy in a discovery cohort (n=10,000), and then the candidate pleiotropic loci were tested for replication in an independent validation cohort of the different genetic ancestry (YRI; n=10,000-35,000). The candidate loci were identified by the JLIM/cFDR consensus method, MetABF and CPBayes. The consensus method (red line), by taking the intersection between JLIM P < 0.01 and cFDR P < 0.01, showed the empirical false positive rate of 0.0038 in simulated $H_0$ dataset. Bayesian posterior thresholds for MetABF (green) and CPBayes (purple) calibrated using $H_0$ loci to match the false positive rate of the consensus method. The 2,500 GWAS peaks consist of the loci simulating no causal effect for underpowered traits ($H_0$) and those simulating the same causal effect between two traits ($H_1$) or distinct causal effects ($H_2$). The proportion of $H_0$ varied to **(A)** 0%, **(B)** 10%, **(C)** 20% and **(D)** 30%, and the remaining loci were split to $H_1$ and $H_2$ at the ratio of 1:19. In all panels, the effect sizes of causative variants are correlated with $\rho$=0.7 under $H_1$ but uncorrelated under $H_2$. Bonferroni correction was applied on replication tests. The shaded area denotes the 95% CIs. In **(E)**, the fraction of $H_1$ among replicated loci is compared among three methods when the proportion of $H_0$ is 30% (similar for different proportions of $H_0$).

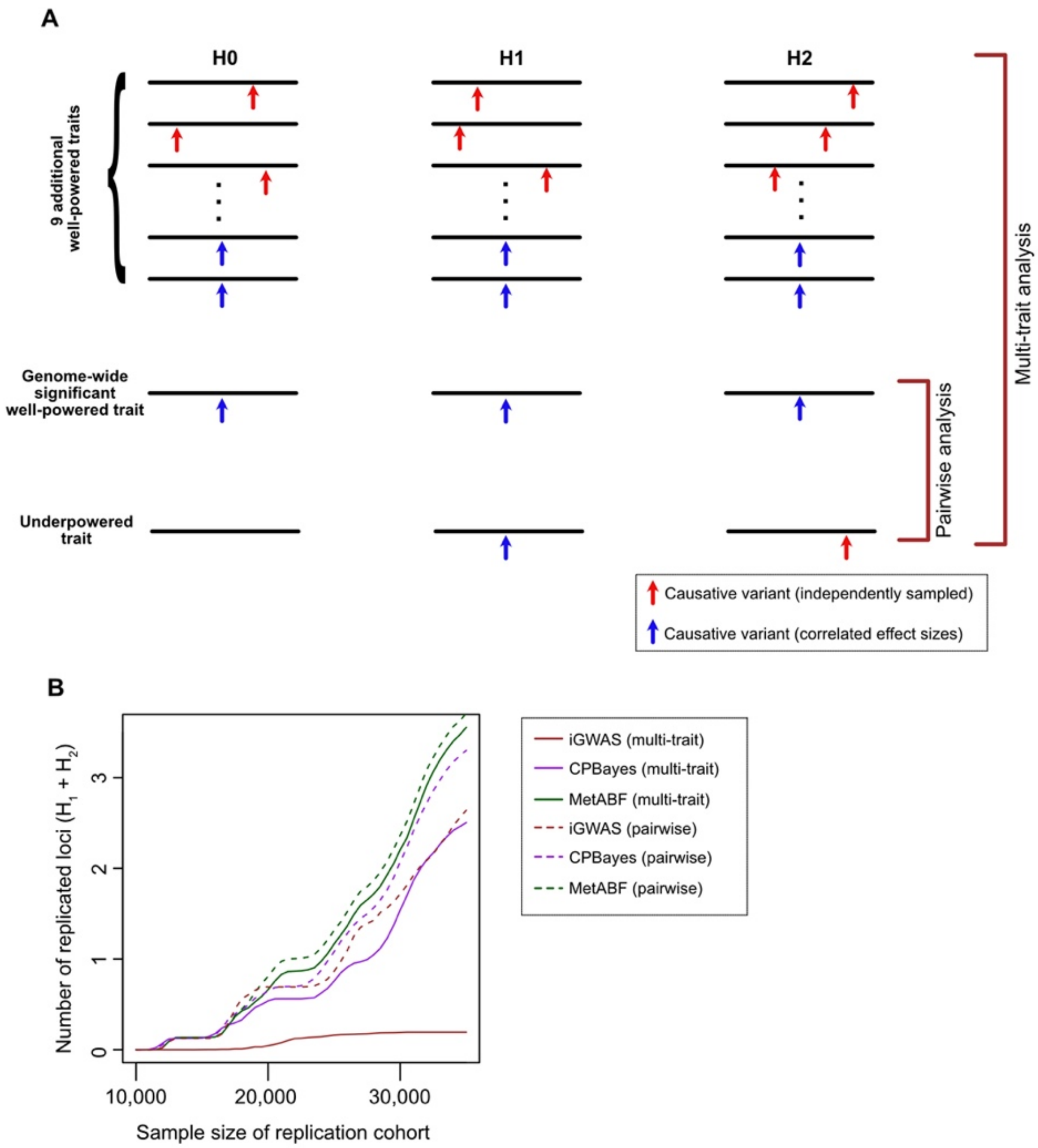


**Figure S15. Simulated power of multi-trait meta-analysis compared to pairwise analysis. (A) Simulation scheme.** Ten well-powered traits and one under-powered trait were simulated under $H_0$, $H_1$ and $H_2$. One of the well-powered traits is ascertained to have a genome-wide significant association peak, and the rest of well-powered traits were randomly decided to have the same or distinct causative variants by sampling from a binomial distribution Binom(n=9, p=1/20). The underpowered trait harbors no causative variant, the same causative variant, or distinct causative variant depending on whether it is simulating $H_0$, $H_1$ or $H_2$. Effect sizes of all traits sharing the same causative variant were sampled together from a multivariate normal distribution with the correlation parameter of 0.7. Effect sizes of traits simulating distinct causative variants were sampled independently. GWAS association statistics at the focal SNP were generated with the sample sizes of n=150,000 for the well-powered traits and n=10,000 for the underpowered trait. The validation cohort of the same ancestry was simulated with the sample sizes of n=10,000 to 35,000. From these simulated $H_0$, $H_1$ and $H_2$ loci, a total of 2,500 GWAS loci were randomly selected at the proportions of 30%, 3.5% and 66.5%, respectively ($H_1$:$H_2$ ratio of 1:19). **(B)** We applied iGWAS, CPBayes and MetABF on these data. Dashed lines indicate pairwise analyses for which only one genome-wide significant well-powered trait was compared with an underpowered trait. Solid lines indicate multi-trait analyses using the full set of ten well-powered traits and one underpowered trait. Candidate pleiotropic loci were selected for replication analysis at the cutoffs of p-value of 0.01 (iGWAS), or equivalent posterior probability thresholds calibrated in the $H_0$ dataset (CPBayes and MetABF). Bonferroni correction was applied on replication tests in the validation cohort.

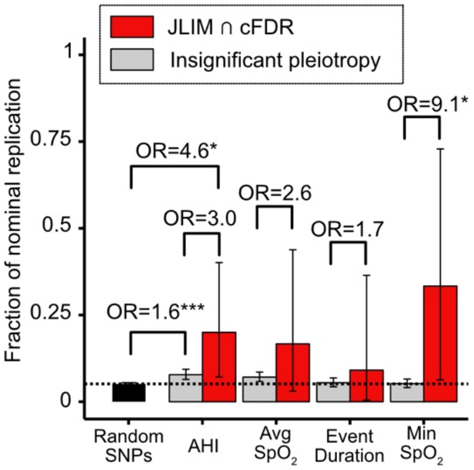


**Figure S16. Pleiotropic loci identified by the intersection of JLIM and cFDR are enriched with SNPs that nominally replicate out of sample.** The plot shows the fractions of randomly selected and putative pleiotropic loci with OSA associations that are nominally replicated in the meta-analyzed independent validation cohort (Table S6).
